# Negligible incorporation of lipophilic dyes into bona fide small extracellular vesicles

**DOI:** 10.64898/2026.07.04.722344

**Authors:** Yuhua Ji, Yuting Shentu, Lingling Zhou, Jingrong Wu, Qian Shao, Mi Shen, Chenhao Zhang, Qiuling Xie, Juling Ji, Qiuhong Ji

**Author notes:** Co-corresponding authors: Juling Ji and Qiuhong Ji. and. **Author information:** These authors contributed equally (Yuhua Ji, Yuting Shentu, Lingling Zhou).

## Abstract

Lipophilic dyes are widely used to track extracellular vesicles (EVs), yet their labeling efficiency toward bona fide small EVs (sEVs) remains poorly defined. Using a serum-free HEK293F system that produces endogenously fluorescent protein–tagged sEVs (sEVs-FPT) as an unambiguous reference while minimizing interference from dye-binding non-vesicular particles, we reassessed this efficiency with two orthogonal methods: nanoflow cytometry and fluorescence microscopy. PKH26, PKH67, and DiD labeled less than 0.5% of sEVs-FPT, regardless of vesicle heterogeneity. In vivo, dye-derived signals were far less abundant than tag-derived signals and failed to colocalize with them. Mechanistic evidence indicates that this failure stems from the inability of sEVs, as acellular structures, to actively internalize dye aggregates. These findings reveal a fundamental limitation of lipophilic dye–based EV tracking and underscore the need for orthogonally validated labeling strategies.

**One Sentence Summary:** Lipophilic dyes fail to label bona fide small extracellular vesicles, calling dye-based vesicle tracking into question.

---

Small extracellular vesicles (sEVs) hold great promise as therapeutic delivery vehicles (*1, 2*), yet realizing this potential critically depends on the ability to reliably track their biodistribution and cellular uptake in vivo (*3, 4*). To date, much of our understanding of EV biodistribution and cellular uptake has been derived from studies using lipophilic fluorescent dyes such as PKH26, PKH67, and DiD, which are favored for their simplicity and strong fluorescence (*5*). These dyes are generally assumed to label sEVs by spontaneous inserting into the lipid bilayer, but their labeling efficiency toward bona fide sEVs remains poorly defined, raising a fundamental question: when we track a dye, are we truly tracking an sEV?

Resolving this question requires a reliable benchmark: a way to unambiguously identify authentic sEVs. Endogenous labeling through genetic expression of a fluorescent protein tag offers near-absolute specificity (*6*). Two pioneering studies adopted this orthogonal approach, using PalmtdTomato- or mEmerald-CD81-tagged EVs as an intrinsic reference against which dye labeling could be assessed (*7, 8*). Intriguingly, both studies found not a complete overlap but a mixed population of dye-positive, tag-positive, and dual-positive particles, with only ∼5 to 20% of tagged EVs being co-labeled by dye. A major confounder likely underlying these puzzling observations is the co-isolation of non-vesicular extracellular particles (NVEPs)—including lipoproteins and protein aggregates—from serum-supplemented culture media. NVEPs overlap with sEVs in size and density and are therefore difficult to remove completely (*9*), yet they readily interact with lipophilic dyes, generating substantial non-sEV fluorescence (*10*). It is thus plausible that a considerable fraction of the dye-positive signals in these studies originated from NVEPs rather than sEVs, obscuring the true labeling efficiency.

To minimize this NVEP interference and critically reassess the reliability of lipophilic dye–based tracking, we established a serum-free HEK293F culture system in which cells secrete sEVs endogenously tagged with fluorescent proteins (sEVs-FPT), providing an unambiguous positive reference with minimal NVEP background. Using two orthogonal and highly sensitive analytical methods—nanoflow cytometry and fluorescence microscopy—we cross-validated the labeling efficiency of commonly used lipophilic dyes against this reference. We show that these dyes label bona fide sEVs with efficiencies below 0.5% and that this deficiency is independent of sEV heterogeneity, persisting across vesicle populations that differ in biogenesis pathway, membrane protein composition, and cellular origin. We further show that dye-derived signals fail to recapitulate the biodistribution and cellular uptake of sEVs in vivo and that this failure stems from a fundamental incompatibility between dye aggregates and the acellular nature of sEVs. These findings resolve the current ambiguity in the field and highlight the need to re-examine how lipophilic dye–based sEV tracking data are interpreted.

## Results

### Generation and validation of genetically labeled sEV reference populations

To evaluate lipophilic dye labeling efficiency without interference from NVEPs, we established a reference system based on serum-free HEK293F cell culture, in which endogenously fluorescent protein–tagged sEVs serve as an internal standard for authentic vesicles. HEK293F cells were transiently transfected with pEGFP-C1-CD63, and CD63-eGFP–tagged sEVs were isolated from the conditioned medium (CM) using the REIUS method (*11*).

Immunoblotting confirmed that the EV markers CD9 and TSG101 were enriched in the concentrated solution (CS), whereas HSP70 was detected in both the CS and the flow-through (FS) (Fig. 1A). CD63-eGFP, detected with both anti-CD63 and anti-GFP antibodies, was predominantly retained in the CS, and fluorescence measurements showed the strongest signal in this fraction (Fig. 1B). After volume normalization, however, more than 50% of total GFP fluorescence was recovered in the FS (Fig. 1C), indicating a substantial pool of soluble or fragmented eGFP in the CM.

**Fig. 1.**
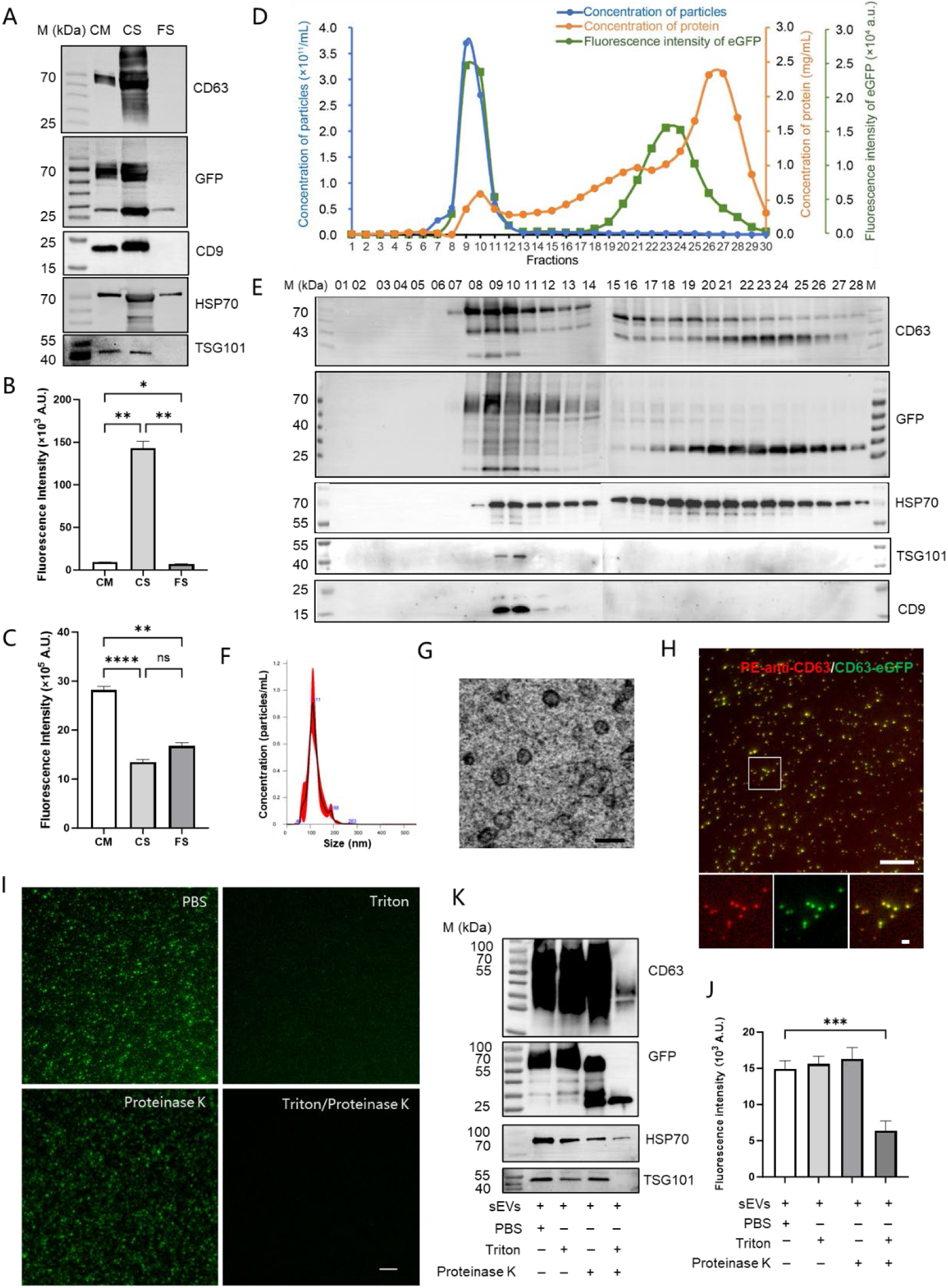
Generation, purification, and characterization of sEVs-CD63-eGFP. (A) Immunoblot analysis of CD63-eGFP and EV markers (CD9, TSG101, HSP70) in conditioned medium (CM), concentrated solution (CS), and flow-through (FS); CS was reconstituted to the original CM volume with PBS before loading. (B and C) Raw (B) and volume-normalized (C) fluorescence intensity of CS and FS. (D) SEC elution profiles of particle concentration, protein concentration, and eGFP fluorescence across 30 fractions. (E) Immunoblot analysis of SEC fractions. (F) NTA size distribution of pooled fractions 9 and 10. (G) TEM image of pooled fractions 9 and 10. Scale bar, 100 nm. (H) Immunofluorescence colocalization of CD63-eGFP (green) with PE-conjugated anti-CD63 antibody (red). Scale bars, 10 μm (overview) and 1 μm (inset). (I to K) Fluorescence images (I), quantification of eGFP intensity (J), and immunoblot analysis (K) after treatment with Triton X-100, proteinase K, or both. Data are presented as mean ± SD, n = 3 independent experiments. \**P* < 0.05, \*\**P* < 0.01, \*\*\**P* < 0.001, \*\*\*\**P* < 0.0001; ns, not significant.

The CS was further fractionated by qEV size-exclusion chromatography (SEC). Approximately 90% of particles eluted in a single peak spanning fractions 6 to 11, which contained ∼40% of total fluorescence but only ∼8% of total protein (Fig. 1D and fig. S1, A to C). CD9 and TSG101 were highly enriched in these fractions, whereas HSP70 showed a broader distribution (Fig. 1E). CD63-eGFP displayed a similar vesicular elution pattern, whereas a ∼25-kDa eGFP-positive band, corresponding to free non-vesicular eGFP, appeared predominantly in later fractions (19 to 27). Fractions 9 and 10, at the center of the vesicle peak, were pooled for downstream analyses. This chromatographic separation of vesicular and free fluorescent protein underscores the need for SEC purification before labeling assessment.

Pooled vesicle-enriched fractions 9 and 10 had a mean diameter of 115 ± 23 nm (mean ± SD) by NTA (Fig. 1F), with no significant size difference from sEVs derived from unmodified cells (fig. S1D), and showed characteristic cup-shaped morphology by TEM (Fig. 1G). Immunostaining showed strong colocalization of anti-CD63 staining with CD63-eGFP fluorescence [Manders’ colocalization coefficient (tM1) = 91.9%], confirming proper membrane localization and topology of the tagged protein (Fig. 1H).

To verify membrane integrity, pooled fractions were treated with Triton X-100 and proteinase K. Whereas Triton X-100 alone disrupted the fluorescent particles, a significant reduction in both fluorescence and protein occurred only with the combined treatment (Fig. 1, I to K), demonstrating that the intact sEV membrane shields luminal CD63-eGFP from proteolysis. Together, these experiments establish sEVs-CD63-eGFP as a membrane-intact reference system for testing lipophilic dye labeling.

### Lipophilic dyes exhibit negligible incorporation into genetically labeled sEVs

We next evaluated lipophilic dye labeling using this reference system. To minimize interference from unbound dye and non-vesicular fluorescent protein, DiD-labeled sEVs-CD63-eGFP were repurified by the REIUS method. Both eGFP and DiD fluorescence were predominantly enriched in the CS (Fig. 2A) and in vesicle-enriched SEC fractions (Fig. 2B). Across all fractions examined, however—CM, CS, FS, and vesicle fractions—the DiD signal remained consistently weak: in the CS and vesicle-enriched fractions, DiD intensity reached only ∼2% of the corresponding eGFP signal after normalization (*P* < 0.0001), quantitatively demonstrating the low labeling efficiency.

**Fig. 2.**
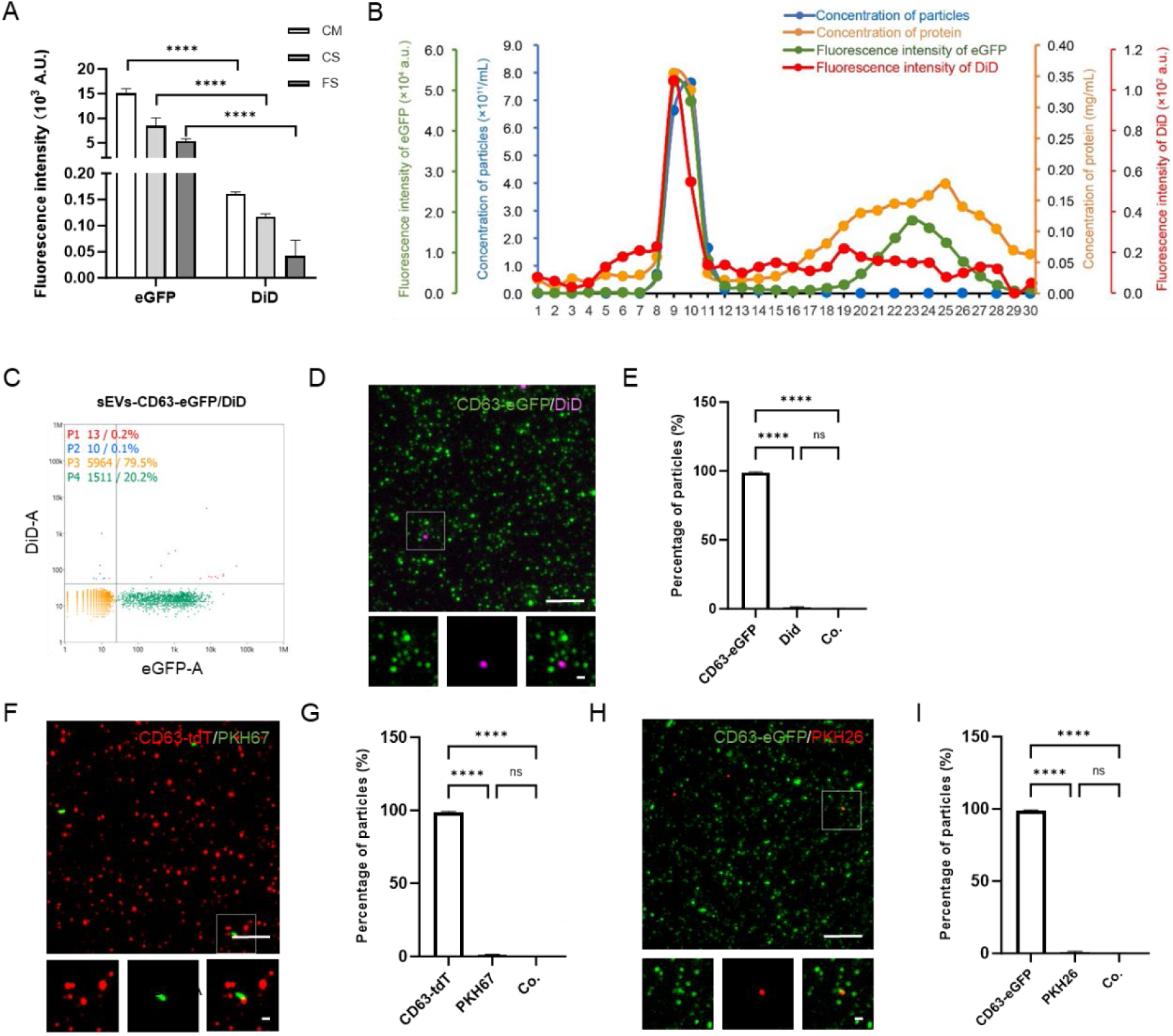
Lipophilic dyes exhibit negligible labeling of sEVs-CD63-FPT. (A) Volume-normalized fluorescence intensity of DiD-labeled sEVs-CD63-eGFP across CM, CS, and FS. (B) SEC co-elution profiles of particle concentration, protein content, and fluorescence across 30 fractions. (C) Representative nFCM dot plot of SEC-purified sEVs-CD63-eGFP labeled with 2 μM DiD. (D) Representative fluorescence micrographs of the same sample. (E) Quantitative image analysis of eGFP⁺, DiD⁺, and dual-positive particles. (F to I) Parallel fluorescence microscopy analyses of sEVs-CD63-tdT labeled with 2 μM PKH67 (F and G) and sEVs-CD63-eGFP labeled with 2 μM PKH26 (H and I). Scale bars, 10 μm (overview) and 1 μm (inset). Data are presented as mean ± SD, n = 3 independent experiments. \*\*\*\**P* < 0.0001; ns, not significant.

Nanoflow cytometry (nFCM) resolved this disparity at the single-particle level. Note that nFCM enumerates all particles by light scattering, whereas fluorescence microscopy detects only fluorescent particles; the particle fractions reported by the two methods therefore have different denominators, with nFCM percentages referring to all particles and microscopy percentages to fluorescent particles. In vesicle-enriched fractions, nFCM detected eGFP⁺, DiD⁺, and dual-positive particles at 20, 0.3, and 0.2% of all particles, respectively (Fig. 2C). Fluorescence microscopy independently confirmed the near-absence of colocalization, revealing abundant, uniformly sized eGFP-positive particles but only a few irregular DiD-positive aggregates—morphologically consistent with co-isolated NVEPs rather than labeled sEVs (Fig. 2D). Among fluorescent particles, >99% were eGFP-positive and <1% were DiD-positive, with virtually no colocalization (Fig. 2E). Increasing the DiD concentration did not increase the DiD-positive fraction (fig. S2, A and B) or alter the particle size distribution (fig. S2C), ruling out insufficient dye concentration as an explanation.

To determine whether this labeling deficiency was specific to DiD or general to lipophilic dyes, we extended the analysis to PKH26 and PKH67 using sEVs-CD63-tdT and sEVs-CD63-eGFP (figs. S3 and S4). All three dyes behaved indistinguishably: PKH26 and PKH67 signals were enriched in the CS and vesicle fractions but remained markedly weaker than the corresponding fluorescent protein signals (*P* < 0.0001; fig. S5), and fluorescence microscopy confirmed that >99% of particles were fluorescent protein–positive, with negligible dye colocalization (Fig. 2, F to I).

Together, these results demonstrate that three widely used lipophilic dyes—DiD, PKH26, and PKH67—achieve negligible labeling of HEK293F-derived sEVs-FPT, with labeling efficiencies consistently below 0.5%.

### sEV heterogeneity does not account for the observed labeling deficiency

Extracellular vesicles are intrinsically heterogeneous, differing in biogenesis pathway, membrane protein composition, and cellular origin (*12*), and such heterogeneity has been proposed to influence lipophilic dye labeling efficiency (*13*). To test whether our findings could be explained by an idiosyncratic feature of CD63-positive vesicles, we systematically evaluated each potential source of variation.

#### Biogenesis pathway

CD63 is primarily associated with exosome-like vesicles originating from multivesicular bodies, whereas CD9 is enriched at the plasma membrane and linked to microvesicle formation (*14, 15*). To assess whether biogenesis origin influences labeling, we generated and characterized sEVs-CD9-eGFP (fig. S6). After DiD labeling, fluorescence microscopy revealed abundant, uniformly sized eGFP⁺ particles alongside a minor population of DiD⁺ structures, with minimal colocalization (Fig. 3, A and B); eGFP⁺, DiD⁺, and dual-positive particles accounted for 98.5, 1.3, and 0.15% of the total, respectively (Fig. 3B). nFCM likewise detected DiD⁺ and dual-positive fractions of only ∼0.2 and ∼0.1% of all particles, respectively (Fig. 3, C and D). Increasing the DiD concentration did not alter this pattern (fig. S7). These results closely paralleled those obtained for CD63⁺ sEVs, indicating that the biogenesis pathway does not explain the labeling deficiency.

**Fig. 3.**
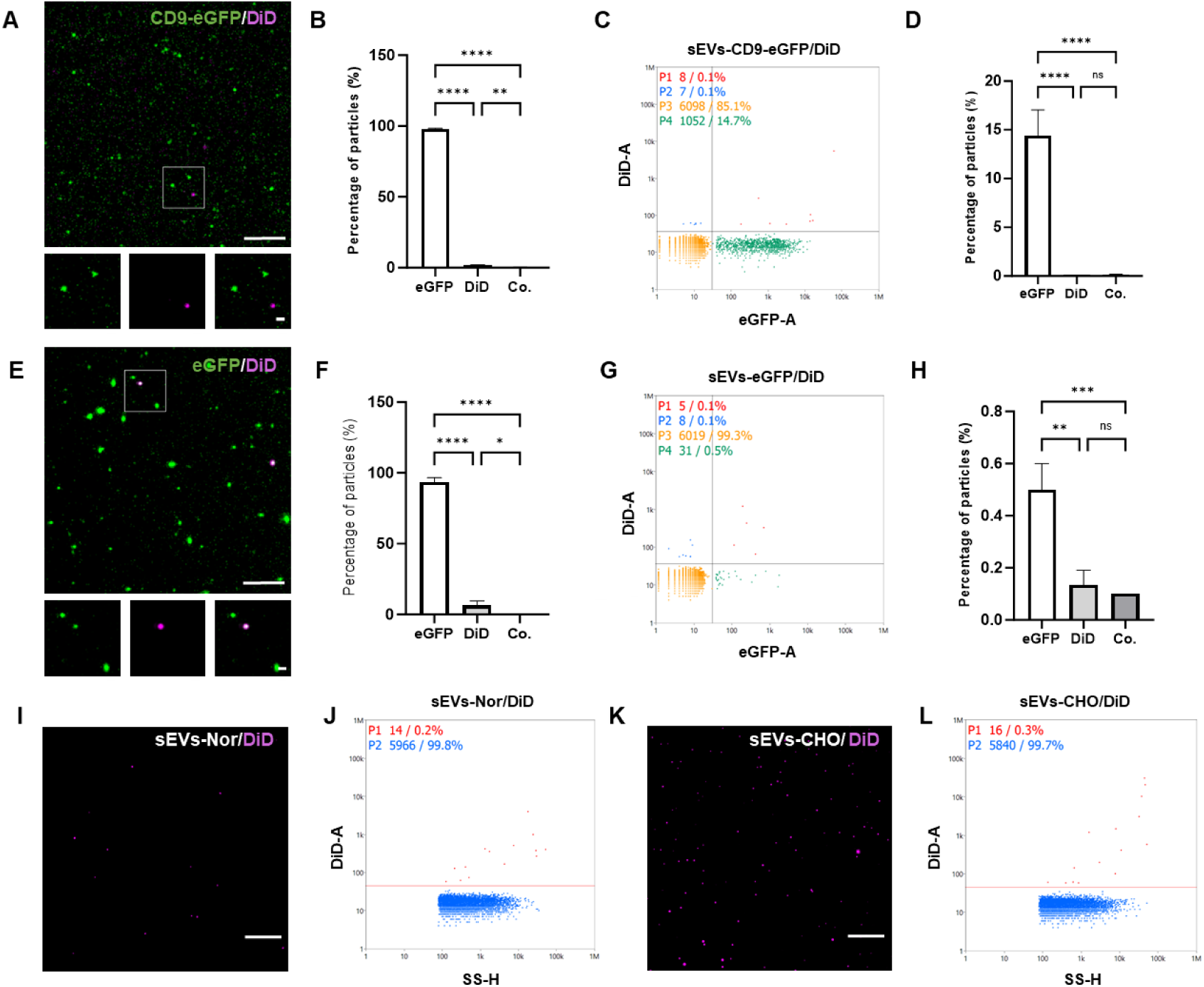
The labeling deficiency is independent of biogenesis pathway, membrane protein composition, and cellular origin. (A to D) DiD-labeled sEVs-CD9-eGFP: representative micrograph (A), quantitative image analysis (B), representative nFCM dot plot (C), and nFCM quantification (D). (E to H) Parallel analysis of DiD-labeled sEVs-eGFP: representative micrograph (E), quantitative image analysis (F), representative nFCM dot plot (G), and nFCM quantification (H). (I and J) DiD-labeled sEVs-Nor: representative micrograph (I) and nFCM dot plot (J). (K and L) DiD-labeled sEVs-CHO: representative micrograph (K) and nFCM dot plot (L). All samples were labeled with 2 μM DiD. Scale bars, 10 μm (overview) and 1 μm (inset). Data are presented as mean ± SD, n = 3 independent experiments. \**P* < 0.05, \*\**P* < 0.01, \*\*\**P* < 0.001, \*\*\*\**P* < 0.0001; ns, not significant.

#### Membrane protein overexpression

We next asked whether overexpression of tetraspanins (CD63 or CD9) could artifactually alter dye labeling. To this end, we generated sEVs-eGFP, which carry soluble eGFP without membrane protein overexpression, and isolated sEVs-Nor from unmodified HEK293F cells (figs. S8 and S9). As previously reported (*16*), sEVs-eGFP contained a markedly smaller eGFP⁺ fraction than CD63- or CD9-tagged sEVs (Fig. 3E and fig. S8B). Despite this difference, DiD labeling again produced predominantly eGFP⁺ particles with negligible colocalization (Fig. 3E), with eGFP⁺, DiD⁺, and dual-positive proportions of 95.5, 4.1, and 0.4% among fluorescent particles (Fig. 3F); nFCM likewise detected DiD⁺ and dual-positive fractions of only ∼0.1% of all particles (Fig. 3, G and H). Increasing the DiD concentration did not alter this pattern (fig. S10). DiD-labeled sEVs-Nor similarly showed only ∼0.2% DiD⁺ particles (Fig. 3, I and J). Thus, tetraspanin overexpression does not artifactually suppress dye labeling.

#### Cellular origin

Finally, we tested whether the labeling deficiency was specific to HEK293F cells. CHO cells, which are adapted to chemically defined medium and widely used in biopharmaceutical production (*17*), provide a distinct cellular background with different membrane composition. DiD labeling of CHO-derived sEVs (sEVs-CHO; fig. S11) produced only a limited number of larger, irregular fluorescent particles (Fig. 3K), with ∼0.3% DiD-positive events by nFCM (Fig. 3L).

Together, these analyses rule out the possibility that the observed labeling deficiency arises from any specific source of EV heterogeneity—biogenesis pathway, membrane protein composition, or cellular origin—under the conditions tested. The inability of lipophilic dyes to efficiently label sEVs therefore appears to be a general phenomenon.

### Lipophilic dyes fail to recapitulate the in vivo biodistribution of sEVs

Lipophilic dyes are widely used for in vivo EV tracking. To assess their reliability in this setting, we intravenously administered REIUS-purified, DiD-labeled sEVs-CD63-eGFP to mice and examined biodistribution and cellular uptake by immunofluorescence across major organs, including the heart, liver, spleen, lung, kidney, and brain.

Among the organs examined, fluorescence signals were detectable only in the liver and spleen (Fig. 4A); neither eGFP nor DiD signals above background were observed in the heart, lung, kidney, and brain (fig. S12), consistent with the known preferential hepatic and splenic accumulation of systemically administered EV preparations (*5*). In these two organs, although the purified vesicle fractions contained only weak DiD fluorescence (Fig. 2B), low-intensity DiD signals remained observable in the tissue sections (Fig. 4A). CD63-eGFP, visualized by anti-GFP immunostaining, was likewise detected in both organs (Fig. 4A). However, the spatial and quantitative profiles of the DiD and CD63-eGFP signals diverged markedly.

**Fig. 4.**
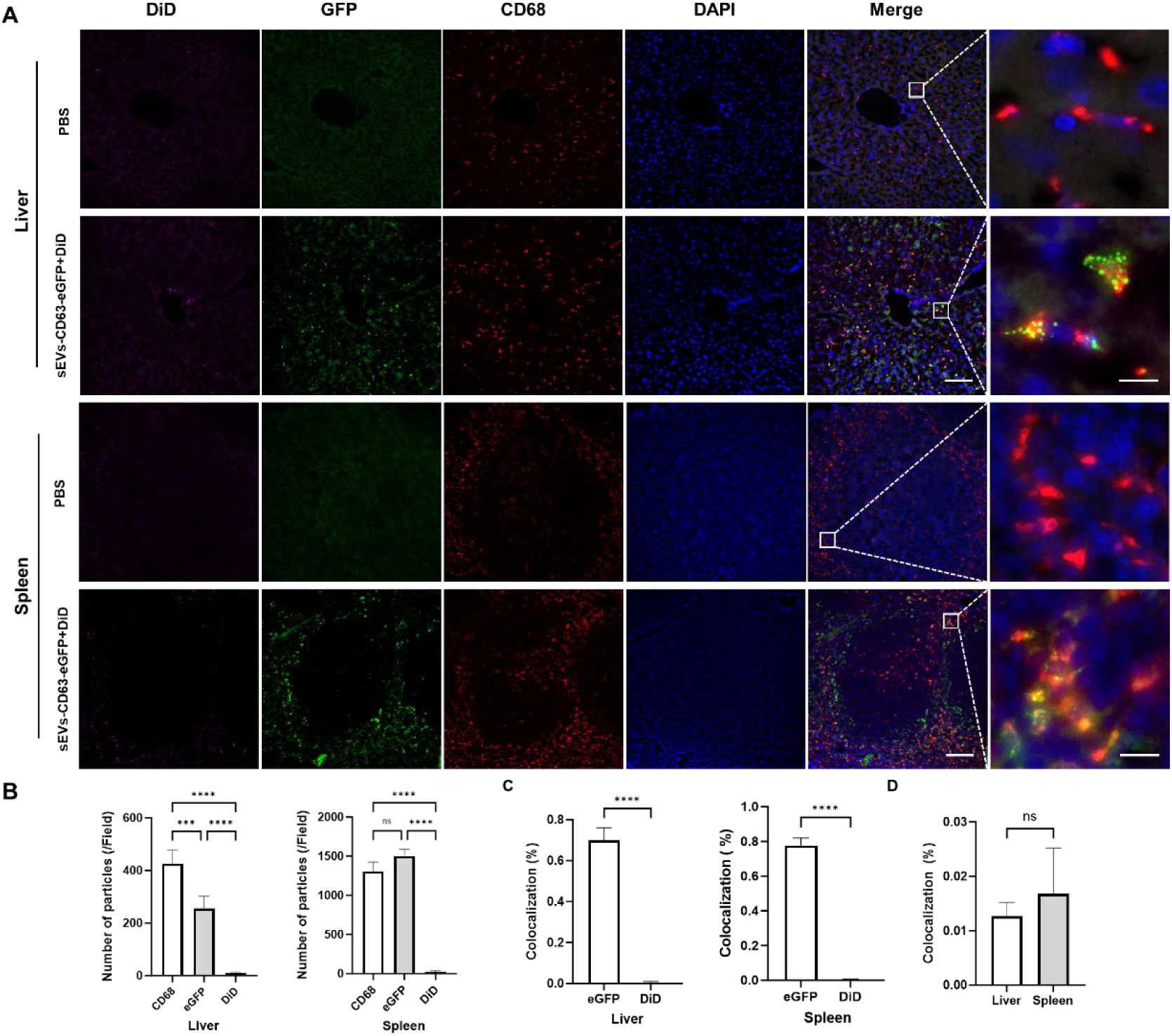
DiD labeling fails to recapitulate the in vivo biodistribution and cellular uptake of sEVs. (A) Representative immunofluorescence images of liver and spleen sections from C57BL/6 mice 30 min after intravenous injection of REIUS-purified, DiD-labeled (2 μM) sEVs-CD63-eGFP; controls received PBS. Sections were stained for eGFP (green) and CD68 (red); DiD (magenta) and DAPI (blue) are shown. No eGFP or DiD signals above background were detected in heart, lung, kidney, or brain sections (fig. S12). (B) Quantification of CD68⁺, eGFP⁺, and DiD⁺ particles in liver and spleen. (C) Colocalization of eGFP and DiD signals with CD68⁺ macrophages. (D) Colocalization between DiD and eGFP signals. Scale bars, 100 μm (overview) and 10 μm (inset). Data are presented as mean ± SD, n = 3 to 4 mice per group. \*\**P* < 0.01, \*\*\**P* < 0.001, \*\*\*\**P* < 0.0001; ns, not significant.

First, eGFP-positive particles outnumbered DiD-positive particles by ∼25-fold in the liver and ∼60-fold in the spleen (*P* < 0.0001; Fig. 4B). Second, their spatial distributions differed fundamentally: DiD fluorescence appeared as sparse puncta, whereas CD63-eGFP showed a vesicular distribution and localized primarily within the cytoplasm of CD68⁺ macrophages in both tissues. Colocalization analysis confirmed substantial overlap between CD63-eGFP and CD68 (tM1 = 69.9% in liver and 75.2% in spleen) but minimal association between DiD and CD68⁺ cells (tM1 < 0.5% in both organs; Fig. 4C). Consistent with our in vitro findings, DiD and CD63-eGFP signals showed virtually no colocalization in vivo (tM1 < 0.05% in both organs; Fig. 4D).

These findings demonstrate that DiD labeling does not faithfully report the biodistribution and cellular uptake of systemically administered sEVs, undermining the reliability of lipophilic dye–based in vivo EV tracking.

### sEVs cannot actively internalize dye aggregates

These findings raise a mechanistic question: why do lipophilic dyes efficiently label HEK293F cells but fail to label their sEVs? Lipophilic dyes readily form self-aggregates in aqueous solution (*18, 19*), and we confirmed this aggregation state in our system. Treatment of DiD- or PKH26-labeled sEVs-CD63-eGFP—analyzed without REIUS repurification so that free dye remained in the mixture—with Triton X-100 or SDS increased fluorescence by nearly two orders of magnitude (Fig. 5, A to D), consistent with aggregation-caused quenching (ACQ), in which fluorescence is suppressed in the aggregated state and restored upon disaggregation (*20*). The dyes therefore exist predominantly as quenched aggregates in the labeling milieu. Notably, the minimal contribution of sEVs to this fluorescence recovery further suggests that the aggregates remain largely free in solution and are not internalized by the vesicles.

**Fig. 5.**
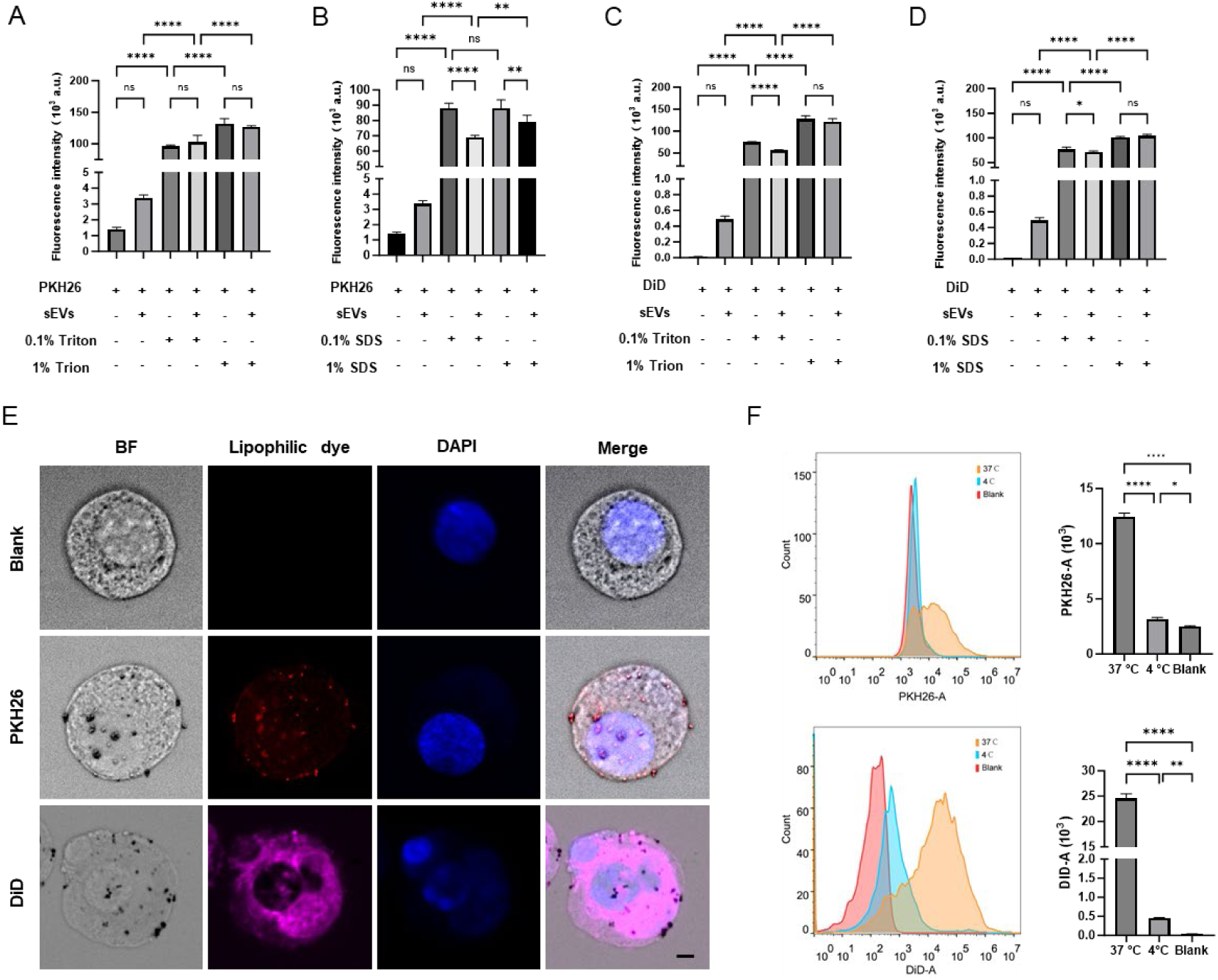
Lipophilic dyes exist as quenched aggregates, and cellular labeling depends predominantly on active internalization. (A to D) Surfactant-induced fluorescence recovery of PKH26-labeled (A and B) and DiD-labeled (C and D) sEVs-CD63-eGFP, analyzed without REIUS repurification so that free dye aggregates remained in the mixture. (E) Representative fluorescence micrographs of HEK293F cells incubated with 2 μM PKH26 or DiD at 37°C. Scale bars, 5 μm. (F) Flow cytometric analysis of HEK293F cells labeled with 2 μM PKH26 or DiD at 37°C or 4°C. Data are presented as mean ± SD, n = 3 independent experiments. \*\**P* < 0.01, \*\*\**P* < 0.001, \*\*\*\**P* < 0.0001; ns, not significant.

Because the labeling process cannot be visualized directly in sEVs with current techniques, we dissected the labeling mechanism in HEK293F cells as a surrogate. Fluorescence microscopy of PKH26- or DiD-labeled cells revealed abundant punctate signals at the plasma membrane and throughout the cytoplasm (Fig. 5E), suggesting that efficient labeling involves active internalization and intracellular disassembly of dye aggregates. To test this directly, we inhibited energy-dependent endocytosis by incubating cells at 4°C. Flow cytometry showed that low temperature reduced DiD and PKH26 fluorescence to ∼2 and ∼25% of control levels, respectively (*P* < 0.001; Fig. 5F), establishing active cellular internalization as the predominant labeling mechanism, with a smaller, dye-dependent contribution from passive membrane insertion reflected in the residual signal at 4°C.

The labeling failure of sEVs can thus be attributed to a fundamental incompatibility: as acellular structures, sEVs lack the endocytic machinery required to internalize and process dye aggregates. The rate-limiting transition from nonfluorescent aggregates to membrane-inserted fluorescent monomers—a step executed efficiently by living cells—cannot be accomplished by these vesicles.

## Discussion

This study was motivated by a fundamental question: when we track a lipophilic dye–labeled particle, are we truly tracking a bona fide sEV? Our findings, obtained in a serum-free system that minimizes the confounding influence of NVEPs, indicate that the answer is no. We demonstrate that three commonly used lipophilic dyes—PKH26, PKH67, and DiD—exhibit negligible labeling efficiency (<0.5%) for sEVs-FPT. This conclusion is supported by two orthogonal analytical methods—nanoflow cytometry and fluorescence microscopy—and is further corroborated by in vivo tracking, in which dye-derived signals were not only markedly less abundant but also failed to colocalize with the FPT signal.

These findings offer a coherent and parsimonious explanation for the heterogeneous particle populations reported by previous orthogonal labeling studies. Both Lai et al. (*7*) and Melling et al. (*8*) described three populations: dye-only, tag-only, and dual-positive particles. Within the framework established by our data, these populations can be rationally reinterpreted: dye-only particles correspond to serum-derived NVEPs that nonspecifically interact with dye aggregates; tag-only particles represent authentic, unlabeled sEVs; and the dual-positive population—previously attributed to successfully labeled sEVs—most likely reflects an artifact in which free fluorescent proteins, which are abundant in conditioned medium even under serum-free conditions (Fig. 1 and figs. S1 and S3), associate with dye-labeled NVEPs. This reinterpretation is consistent with the discrepancy in the reported dual-positive rates—∼20% in the ultracentrifugation (UC)-based study (*7*) versus ∼5% in the SEC-based study (*8*)—because UC-purified EV fractions contain substantially more co-isolated soluble protein than SEC-purified fractions, with protein content differing by up to 45-fold (*21, 22*).

Our data also provide insight into the mechanistic basis of this labeling failure. The fluorescence of lipophilic dyes depends on a critical transition from nonfluorescent aggregates to membrane-inserted monomers. In HEK293F cells, labeling was strongly dependent on active endocytosis—which internalizes dye aggregates—and was largely abolished at low temperature, whereas a passive, membrane fluidity–dependent diffusion pathway likely makes a smaller contribution. As acellular nanostructures, sEVs lack the endocytic machinery required to internalize and process dye aggregates. Moreover, the hydrophilic protein corona present on the sEV surface (*23*) may pose an additional physical barrier, limiting both the contact and passive fusion of hydrophobic dye aggregates with the vesicle membrane. Together, these factors restrict the rate-limiting monomerization step on sEVs. This aggregate-to-monomer bottleneck is consistent with a recent report that chemical disruption of dye aggregates through ionic modulation markedly improves vesicle labeling (*24*).

This mechanistic model is based primarily on HEK293F cells. Because endocytic capacity and membrane dynamics can vary across cell types, its generalizability awaits further investigation. Nevertheless, the core principle—that an acellular vesicle cannot actively process particulate dye aggregates—is likely a universal constraint for any nanoparticle tracking strategy that relies on a similar aggregate-to-monomer transition. Elucidating how living cells accomplish this transition—and why vesicles cannot—will guide the development of more efficient labeling methods.

The implications of our study extend beyond lipophilic dyes. A wide range of post-labeling strategies—including the conjugation of fluorophores, radionuclides, or nanoparticles to surface proteins or nucleic acids (*25*)—share the same fundamental vulnerability to co-labeling of non-vesicular contaminants. Free proteins and nucleic acids present in any biofluid or conditioned medium can act as unintended acceptors of labeling probes and thereby confound tracking analyses. Where post-labeling is unavoidable, as in clinical samples, rigorous and context-specific validation of labeling specificity is not merely advisable but imperative.

In accordance with the MISEV2023 guidelines (*10*) and building on the present findings, we propose a four-pillar framework for rigorously validating post-labeling efficiency for sEVs, resting on four pillars: (1) an independently labeled, unambiguous sEV reference must be established, for which endogenous fluorescent protein tagging currently provides the most rigorous approach; (2) isolation procedures, such as SEC, must be employed to minimize confounding by unbound labeling reagents and co-isolated non-vesicular contaminants, including free fluorescent proteins; (3) orthogonal analytical techniques should be combined to reduce methodological bias while providing both qualitative and quantitative readouts, as demonstrated here with fluorescence microscopy and nanoflow cytometry; and (4) where possible, in vivo tracking experiments should be conducted to confirm that labeling specificity is maintained in the complex physiological environment.

In summary, our study reveals that lipophilic dyes—the most widely used tools for EV tracking—fail to label bona fide sEVs under serum-free conditions, with labeling efficiencies below 0.5%. This failure stems from the inability of acellular sEVs to convert dye aggregates into fluorescent monomers, a step that in living cells is accomplished through active endocytosis. These findings demonstrate that lipophilic dye labeling cannot be assumed to report bona fide sEVs; conclusions drawn solely from dye-derived signals therefore await orthogonal confirmation. Rigorous, independently validated labeling frameworks will be essential for future sEV tracking studies.

## Materials and Methods

### Cell culture and transfection

HEK293F cells (Expi293F, Thermo Fisher, Cat# A14527) were cultured in chemically defined OPM-293 CD05 medium (OPM Biosciences, Cat# 81075-001) in 500-mL Erlenmeyer flasks (Jetbiofil) at 37 °C, 8% CO₂, and 125 rpm (ZCZY-ANS shaker, ZHICHU Biotech). Human CD63 (UniProt P08962) and CD9 (UniProt P21926) coding sequences were synthesized (Sangon Biotech) and cloned into pEGFP-C1 or ptdTomato-C1 vectors (IBSBIO), and all plasmids were verified by sequencing. HEK293F cells were transiently transfected using PEI MAX (40 kDa, 1 mg/mL; Polysciences) at a DNA dose of 1 μg per 10⁶ cells (PEI:DNA ratio of 2:1). Cell viability at 48 h post-transfection consistently exceeded 90%, as determined by trypan blue exclusion (CountStar, Alit Biotech). Conditioned medium (CM) was sequentially centrifuged at 300 × g for 10 min (cells), 2,000 × g for 10 min (cell debris), and 10,000 × g for 30 min (large particles), all at 4 °C, and the final supernatant was passed through a 0.22-μm filter (Millipore).

CHO-S cells (Thermo Fisher, Cat# A29127) were cultured in ProCHO-5 medium supplemented with 4 mM glutamine, 0.68 mg/L hypoxanthine, and 0.194 mg/L thymidine (Lonza) in TubeSpin Bioreactor 50 tubes (TPP) at 37 °C, 8% CO₂, and 75–90 rpm. CM was collected following the same procedure.

### REIUS isolation and purification of sEVs

sEVs were isolated and purified using the REIUS method (*11*). Briefly, prefiltered CM was concentrated with Amicon Ultra-15 100-kDa molecular weight cut-off (MWCO) centrifugal filters (Merck Millipore, UFC810024), and both the concentrated solution (CS) and the flow-through solution (FS) were collected. Size-exclusion chromatography (SEC) was performed on a qEV original 35-nm column (Izon Science), which was washed with 0.1-μm-filtered PBS (Biosharp, BL302A) and equilibrated at room temperature. A 0.5-mL aliquot of concentrated

CM was loaded, eluted with PBS, and 30 fractions (0.4 mL each) were collected. Vesicle-enriched fractions (fractions 9 and 10) were pooled and, if necessary, further concentrated using 100-kDa MWCO centrifugal filters. Final sEV samples were aliquoted and stored at −80 °C.

### Nanoparticle tracking analysis

Particle size and concentration were measured by nanoparticle tracking analysis (NTA) using a NanoSight NS300 instrument equipped with a 488-nm laser (NTA software v3.3; Malvern Panalytical). Samples were diluted in 0.1-μm-filtered PBS to approximately 1 × 10⁸–1 × 10⁹ particles/mL. Measurements were carried out at 25 ± 0.01 °C with continuous syringe pump infusion. For each sample, three 30-s videos were captured. The camera level (11–13), screen gain (10–12), and detection threshold (3–5) were maintained within the indicated ranges, with identical settings applied to all samples within each experiment. Videos were analyzed with NTA software to determine mean particle size, size distribution, and particle concentration.

### Fluorescence measurement and protein quantification

Fluorescence intensities of pre- and post-ultrafiltration samples and SEC fractions were measured using a Synergy H1 multimode microplate reader (BioTek Instruments) with Gen5 software (v2.09) in monochromator mode. All readings were taken in top-read mode with the detector gain fixed at 100, and the detection range was held identical across channels, enabling direct comparison of signal intensities between channels. Excitation/emission wavelengths were 554/581 nm (tdTomato, PKH26), 488/502 nm (eGFP, PKH67), and 644/665 nm (DiD). Blank controls (PBS or the corresponding buffer) were included for background subtraction. Each sample was measured in triplicate, and fluorescence intensity was expressed as relative fluorescence units (RFU). Protein concentrations of SEC fractions were quantified using the Pierce BCA Protein Assay Kit (Thermo Fisher Scientific, Cat# 23227) according to the manufacturer’s instructions.

### Western blotting

Samples were mixed with 5× SDS-PAGE loading buffer (Beyotime, Cat# P0015L) containing DTT and denatured at 90 °C for 15 min. Proteins were separated by SDS-PAGE and transferred onto PVDF membranes (Millipore). Membranes were blocked with blocking buffer (Beyotime, Cat# P0951) at room temperature and then incubated overnight at 4 °C with primary antibodies diluted in blocking buffer: anti-eGFP (Abcam, Cat# ab6673, 1:1000), anti-CD63 (Abcam, Cat# ab59479, clone TS63, 1:1000), anti-CD9 (Proteintech, Cat# 25682-1-AP, 1:1000), anti-HSP70 (Proteintech, Cat# 10995-1-AP, 1:10,000), anti-TSG101 (Proteintech, Cat# 28283-1-AP, 1:3000), and anti-tdTomato (BBI, Cat# D199981-0100, 1:3000). After three washes with TBST, membranes were incubated for 1 h at room temperature with HRP-conjugated secondary antibodies diluted 1:2000 in TBST: goat anti-rabbit IgG H&L (Beyotime, Cat# A0208), goat anti-rat IgG H&L (Beyotime, Cat# A0192), and goat anti-mouse IgG H&L (Beyotime, Cat# A0216). After additional TBST washes, bands were visualized using BeyoECL Plus (Beyotime, Cat# P0018S) and imaged with a Tanon chemiluminescence imaging system (Tanon).

### Transmission electron microscopy

A 20-μL aliquot of purified sEV suspension (∼1 × 10¹¹ particles/mL) was adsorbed onto a 200-mesh Formvar/carbon-coated copper grid for 3 min at room temperature. Excess liquid was blotted away with filter paper, and the grid was rinsed with Milli-Q water to remove buffer salts. Samples were negatively stained with 1% (w/v) uranyl acetate for 30 s, air-dried at room temperature, and examined using a JEOL JEM-1400 Flash transmission electron microscope (JEOL).

### Immunofluorescent staining of sEVs

Purified sEVs-CD63-eGFP (10 μL) were incubated with PE-conjugated anti-CD63 antibody (BioLegend, Cat# 353004; 1:100) for 15 min at 4 °C in the dark. A PE-conjugated isotype control (BioLegend, Cat# 400111) was used to assess nonspecific binding.

### Vesicle protection assay

The vesicle protection assay was performed as described (*26*) with modifications. Purified sEVs typically had a protein concentration of 200–300 μg/mL (BCA assay). Proteinase K was used at an enzyme-to-protein ratio of 1:50, yielding a final concentration of 5 μg/mL. Triton X-100 was used at a final concentration of 0.1% (v/v), which was confirmed to permeabilize sEV membranes under the described conditions. Purified sEVs (100 μL) were subjected to one of the following treatments: (i) 0.1% Triton X-100 (Sigma-Aldrich, Cat# T8787), (ii) 5 μg/mL proteinase K (Roche, Cat# 03115879001), (iii) 0.1% Triton X-100 plus 5 μg/mL proteinase K, or (iv) PBS (untreated control). All reactions were incubated at 37 °C for 15 min. Following treatment, each sample was divided into three aliquots for Western blotting, fluorescence quantification by microplate reader, and fluorescence microscopy imaging.

### Lipophilic dye labeling of sEVs

sEVs were labeled with lipophilic dyes according to the manufacturers’ instructions with modifications. For PKH labeling, purified sEVs (1 × 10¹¹ particles/mL) were mixed with PKH26 (MCE, Cat# HY-D1451) or PKH67 (Sigma-Aldrich, Cat# MINI67) prepared in Diluent C; for DiD labeling, sEVs at the same concentration in PBS were mixed with DiD (Beyotime, Cat# C1039). Final dye concentrations ranged from 0.1 to 10 μM, depending on the experiment. Labeling reactions were incubated in the dark at room temperature for 10 min. The conventional quenching step using FBS or BSA was omitted because preliminary experiments showed that BSA itself can be labeled, forming non-vesicular fluorescent particles, consistent with recent findings (*27*). Labeled sEVs were subsequently purified by the REIUS method to remove unbound dye.

### Imaging and quantitative analysis of sEVs

Imaging and quantification were adapted from published methods with minor modifications (*7, 28*). Purified or labeled sEVs (10 μL) were mixed with an equal volume of anti-fade mounting medium (Beyotime, Cat# P0126), and 2 μL of the mixture was mounted onto a glass slide and coverslipped. Fluorescence images were acquired on a Leica THUNDER Imager with a 63×/1.4 NA oil-immersion objective, using identical acquisition parameters (laser intensity, exposure time, gain, and threshold) across all groups. Image processing was performed with LAS X software. Particle quantification was carried out using the Analyze Particles plugin in ImageJ (v1.54m) (*29*). Colocalization was analyzed with the Coloc 2 plugin, using identical thresholds and processing settings across all samples.

### Nanoflow cytometry analysis

sEV samples were analyzed by nanoflow cytometry (nFCM) using a NanoFCM instrument (NanoFCM Inc.) equipped with 488- and 638-nm lasers. All measurements were performed according to the manufacturer’s instructions, with distilled water as the sheath fluid. The instrument was calibrated for particle size using a mixture of monodisperse silica nanoparticles with defined diameters (53, 73, 96, and 120 nm; S23M SEV, NanoFCM Inc.), and fluorescence calibration was performed using QC fluorescent microspheres (QS2503, NanoFCM Inc.) to standardize detector sensitivity across the FITC and PC5 channels. All calibration procedures used the same optical and fluidic settings as the experimental samples.

Laser alignment and instrument performance were optimized daily to achieve maximal signal intensity and stability in the FITC, PC5, and side scatter (SSC) channels. The detection threshold was configured using the manufacturer’s ultra-small mode to enhance sensitivity for nanoscale particle detection. Event triggering was based on SSC signals.

Samples were diluted in 0.1-μm-filtered PBS to approximately 1 × 10⁹ particles/mL, yielding event rates within the recommended range of 2,000–12,000 particles per minute. Each sample was acquired for 1 min at a sampling pressure of 1.0 kPa. Filtered PBS served as a negative control to define background noise, and the threshold was set so that the PBS control count was below 10 events per minute, ensuring that signals above this threshold in sEV samples represented bona fide particles.

Detergent lysis controls were performed to verify the vesicular nature of detected particles: sEV aliquots were treated with 0.1% (v/v) Triton X-100 for 10 min at room temperature, and a marked reduction in particle counts after treatment confirmed that the detected signals originated from membrane-enclosed vesicles rather than protein aggregates or other non-vesicular contaminants.

Data acquisition and analysis were performed using NanoFCM Professional software (v3.0). Signal display parameters were uniformly adjusted across all samples to optimize visualization of low-intensity nanoparticle events while maintaining consistency between experimental groups.

### Surfactant-induced dequenching of dye aggregates

Labeled sEV samples were analyzed without REIUS repurification, such that unbound dye aggregates remained in the mixture. Samples were treated with 0.1% (v/v) Triton X-100 or 0.1% (v/v) SDS for 5 min at room temperature, and fluorescence was measured before and after treatment on the Synergy H1 microplate reader using the channel settings described above. Fluorescence recovery was expressed as the fold change relative to untreated controls.

### Lipophilic dye labeling of HEK293F cells

Labeling was performed according to the manufacturer’s protocol. Exponentially growing HEK293F cells were washed twice with PBS. For PKH26 labeling, cells were resuspended in Diluent C (2 × 10⁶ cells/mL); for DiD labeling, cells were resuspended in PBS (2 × 10⁶ cells/mL). The cell suspension was divided into two equal aliquots and pre-equilibrated at 37 or 4 °C for 1 h. PKH26 (MCE, Cat# HY-D1451) or DiD (Beyotime, Cat# C1039) was then added to a final concentration of 2 μM, and cells were incubated at the respective temperatures for an additional 30 min. Unbound dye was removed by centrifugation at 4 °C, and cells were resuspended in PBS. Images were acquired using a Leica THUNDER Imager and processed with LAS X software. Cellular fluorescence was measured by flow cytometry (DxFLEX, Beckman Coulter) and analyzed with FlowJo X.

### In vivo biodistribution of systemically administered sEVs

Male C57BL/6J mice (12–14 weeks, 22–25 g; Huachuang Sino) were housed under specific pathogen-free conditions with a 12-h light/dark cycle and free access to food and water. All procedures followed NIH guidelines and were approved by the Ethics Committee of the Affiliated Hospital of Nantong University (approval no. 2020-L085).

For biodistribution studies, anesthetized mice (inhaled isoflurane; Shenzhen Ruiwode Life Technology) received retro-orbital injections of DiD-labeled sEVs (5 × 10¹⁰ particles per mouse; n = 3–4 mice per group) or an equal volume of PBS (control group, n = 2–3 mice per group). Mice were randomly assigned to experimental groups. Thirty minutes after injection, mice were deeply anesthetized and transcardially perfused with 30 mL of PBS followed by 20 mL of 4% paraformaldehyde (PFA) to clear blood and minimize background. The brain, heart, liver, spleen, lungs, and kidneys were harvested and post-fixed in 4% PFA at 4 °C for 24 h.

### Immunofluorescent staining and image analysis

After fixation, tissues were dehydrated in 10, 20, and 30% sucrose at 4 °C, embedded in optimal cutting temperature (OCT) compound (Tissue-Tek, Sakura), frozen, and sectioned at 8 μm using a cryostat (Leica CM1950). Sections were mounted on poly-L-lysine–coated slides and stored at −20 °C. For immunofluorescence, sections were equilibrated to room temperature, rinsed with PBS, and blocked with blocking buffer (Beyotime, Cat# P0260) for 3–4 h in a humidified chamber. For simultaneous detection of eGFP and CD68, sections were incubated overnight at 4 °C with rat anti-EGFP/EYFP polyclonal antibody (Oasis Biofarm, Cat# OB-PRT018; 1:200) and rabbit anti-CD68 antibody (Boster, Cat# BA3638; 1:200). After washing with PBS, sections were incubated for 1 h at room temperature in the dark with FITC-conjugated goat anti-rat IgG H&L (Abcam, Cat# ab6840; 1:1000) and Alexa Fluor 555–conjugated donkey anti-rabbit IgG H&L (Abcam, Cat# ab150073; 1:1000). After final PBS washes, slides were mounted with anti-fade mounting medium containing DAPI (Beyotime, Cat# P0131).

Fluorescence images were acquired on a Leica THUNDER Imager under identical exposure settings and processed with LAS X software. Quantitative analysis was performed using ImageJ (v1.54m). Positive signals were quantified with the Analyze Particles plugin after background subtraction and threshold segmentation, and colocalization was analyzed with the Coloc 2 plugin. For each tissue, 3–5 sections were analyzed per animal, with at least four fields imaged per section. Thresholds and analysis parameters were established on negative control sections and applied uniformly across all experimental groups.

### Statistical analysis

Data are presented as means ± SD. Statistical analyses were performed in GraphPad Prism 7.05. Comparisons between two groups were made using two-tailed Student’s *t* tests; comparisons among three or more groups used one-way ANOVA followed by Tukey’s or Dunnett’s post hoc test. *P* < 0.05 was considered statistically significant. Unless otherwise noted, all data are representative of at least three independent experiments, and the exact *n* for each experiment is reported in the corresponding figure legend. Mice were randomly assigned to experimental groups; investigators were not blinded to group allocation during data collection and analysis.

## Data and materials availability

All data needed to evaluate the conclusions in the paper are present in the paper or the supplementary materials. Plasmids generated in this study are available from the corresponding author upon request.

## Author contributions

Yuhua Ji: Conceptualization, Funding acquisition, Writing – original draft, Writing – review & editing. Yuting Shentu: Investigation, Formal analysis, Visualization, Writing – original draft. Lingling Zhou: Investigation, Formal analysis, Writing – original draft, Visualization, Validation. Jingrong Wu: Investigation, Formal analysis. Qian Shao: Validation, Data curation. Chenhao Zhang: Formal analysis, Visualization. Mi Shen: Methodology, Resources. Qiuling Xie: Methodology, Resources. Juling Ji: Conceptualization, Supervision, Funding acquisition, Writing – review & editing. Qiuhong Ji: Conceptualization, Supervision, Project administration, Funding acquisition, Writing – review & editing. All authors read and approved the final manuscript.

## Acknowledgment

The authors thank Dr. Xinmei Yu, Application Specialist at NanoFCM Inc., for her technical support in sample analysis and data processing.

