## Supplementary material for "Negligible incorporation of lipophilic dyes into bona fide small extracellular vesicles": Supplemtal figures

### Supplementary Figures

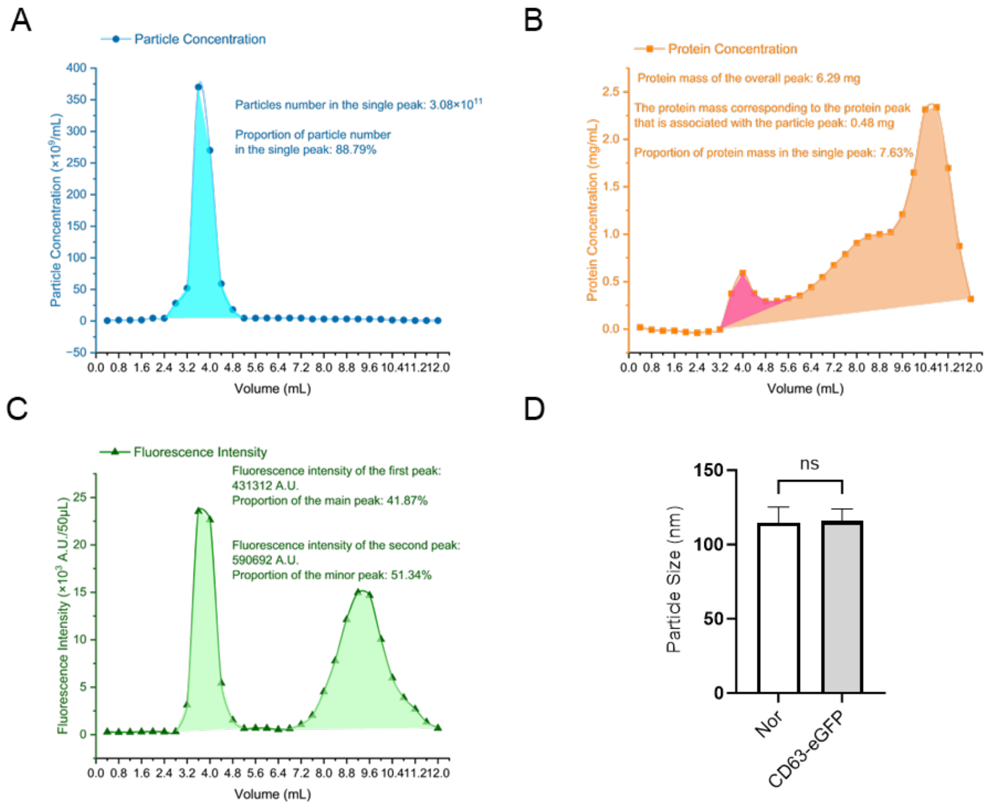


**fig. S1. Quantitative profiling of SEC fractions for sEVs-CD63-eGFP purification.** Raw data for particle number, protein concentration, and eGFP fluorescence intensity across SEC fractions were analyzed and fitted using Origin (v2025sr1). (A) Particle number distribution fitted to a single-peak model. The major peak (fractions 6–11, 2.4–4.4 mL) contained >88% of total particles. (B) Protein concentration profile fitted to a multi-peak model. Particle-rich fractions (6–11) accounted for only 7.63% of total protein, indicating efficient separation from soluble contaminants. (C) eGFP fluorescence intensity fitted to a multi-peak model, revealing a bimodal distribution; the primary peak co-eluting with particles (fractions 6–11) represented 41.87% of total fluorescence. (D) Comparison of particle diameter between sEVs from unmodified (sEVs-Nor) and CD63-eGFP–overexpressing HEK293F cells (sEVs-CD63-eGFP), showing no significant difference in vesicle size. Data are presented as mean ± SD; n = 3 independent experiments. ns, not significant.

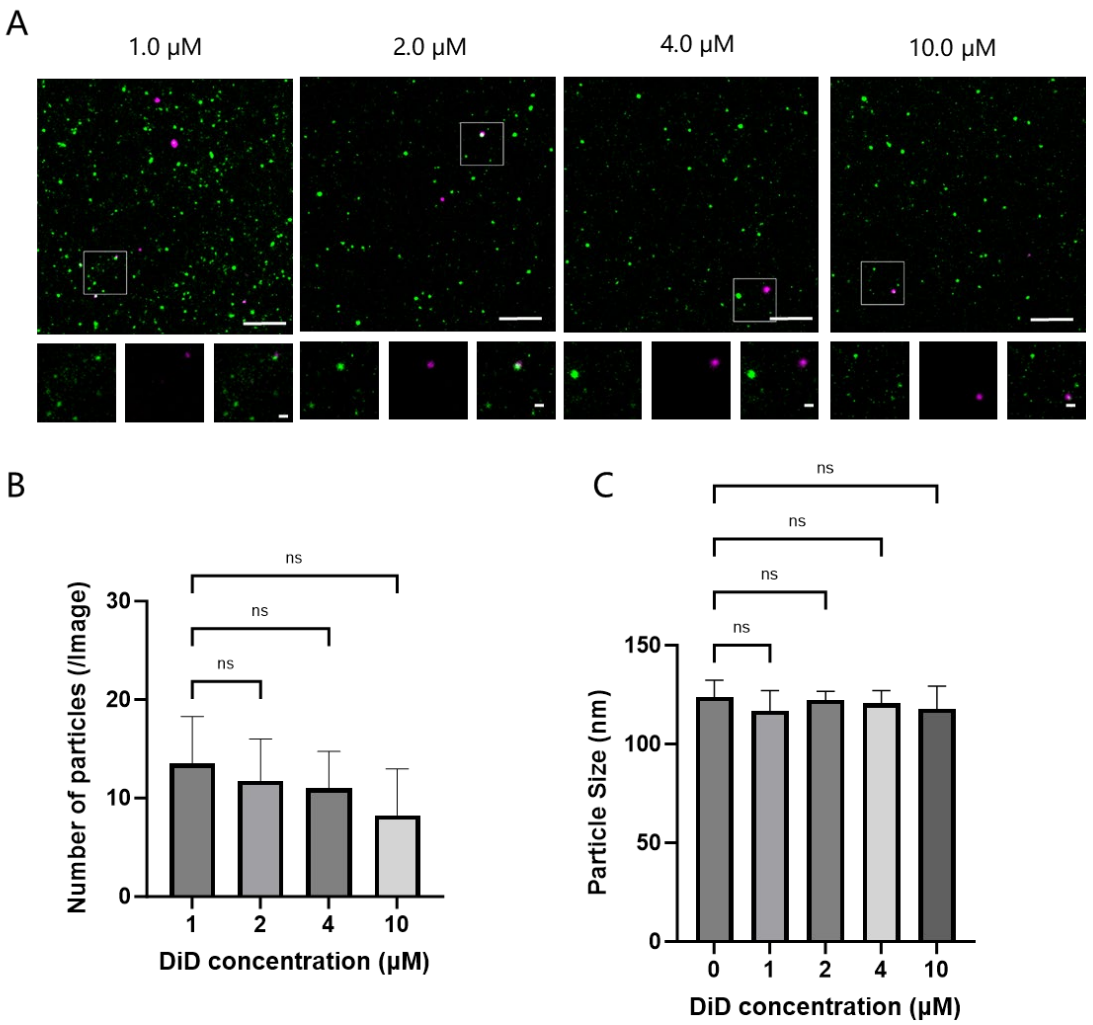


**fig. S2. Concentration-dependent assessment of DiD labeling on sEVs-CD63-eGFP.** (A) Representative fluorescence micrographs of sEVs-CD63-eGFP labeled with increasing concentrations of DiD (1, 2, 4, and 10 μM). (B) Quantitative image analysis of DiD-positive particle counts across the tested concentration range. (C) NTA analysis showing that DiD labeling did not alter the size distribution of sEVs compared with the unlabeled control. Scale bars, 10 μm (overview) and 1 μm (inset). Data are presented as mean ± SD; n = 3 independent experiments. ns, not significant.

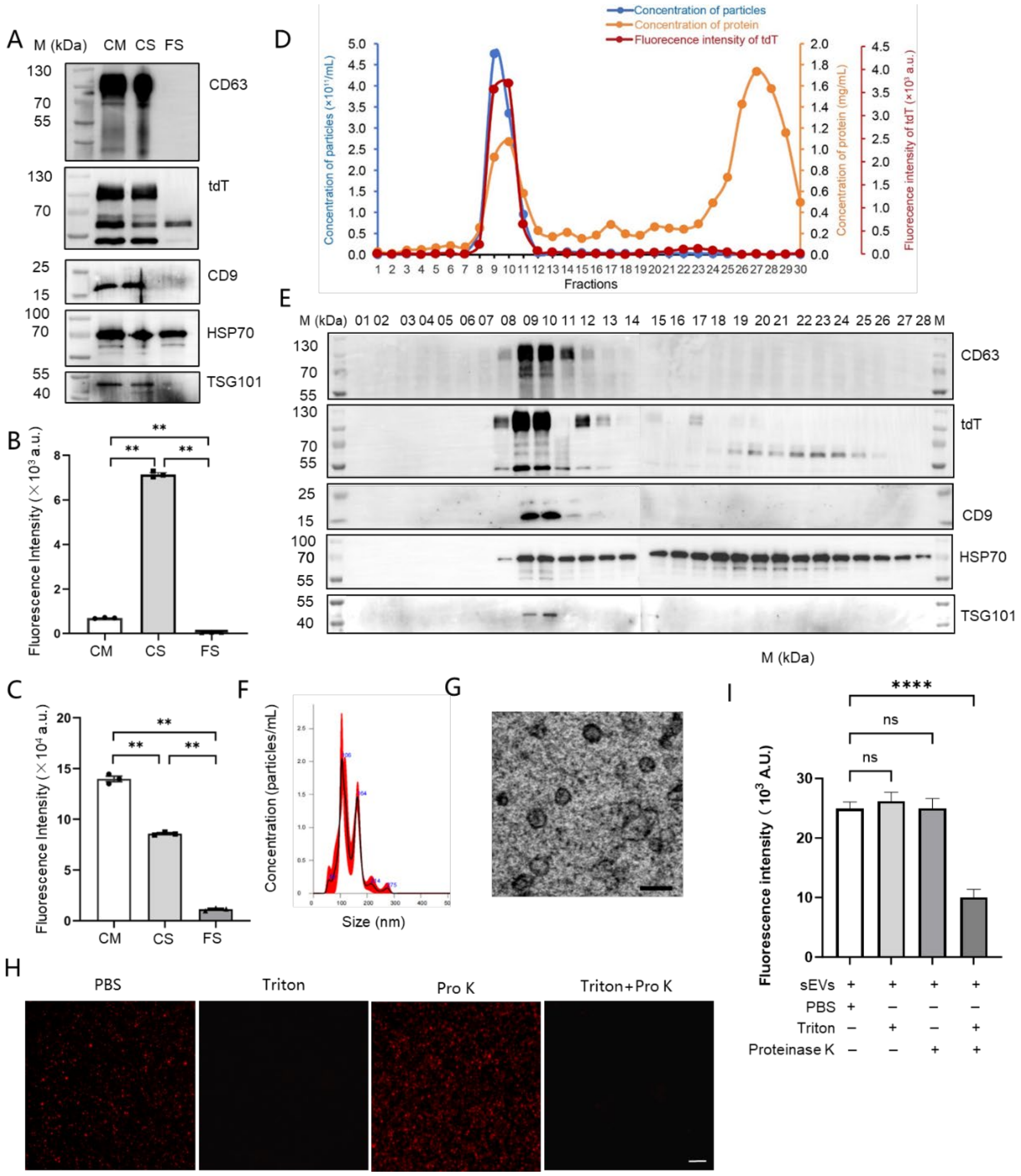


**fig. S3. Generation, purification, and characterization of sEVs-CD63-tdT.** sEVs were purified from the conditioned medium (CM) of HEK293F cells transiently transfected with ptdTomato-C1-CD63 using the REIUS method. (A) Immunoblot analysis of CD63-tdT and EV markers (CD9, TSG101, HSP70) in CM, concentrated solution (CS), and flow-through (FS); CS was volume-adjusted with PBS for quantitative comparison. (B and C) Raw (B) and volume-normalized (C) tdTomato fluorescence intensity before and after ultrafiltration. (D) SEC co-elution profiles of particle concentration, protein concentration, and tdTomato fluorescence. (E) Immunoblot analysis of CD63-tdT and EV markers (CD9, TSG101, HSP70) in SEC fractions. (F) NTA size distribution of pooled fractions 9 and 10. (G) TEM image confirming vesicular morphology. Scale bar, 100 nm. (H and I) Membrane integrity assessment by fluorescence microscopy (H) and quantitative analysis (I) of tdTomato signal after treatment with Triton X-100, proteinase K, or both. Scale bars, 10 μm. Data are presented as mean ± SD; n = 3 independent experiments. *$P < 0.01$, ***$P < 0.0001$; ns, not significant.

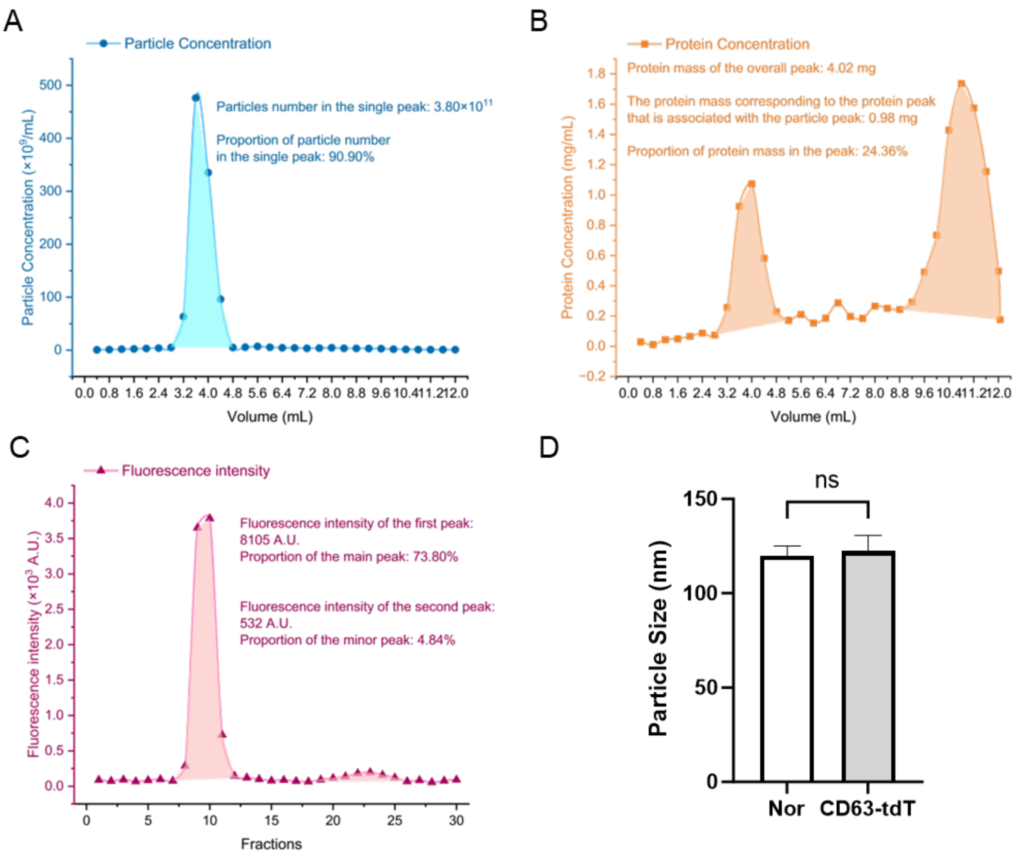


**fig. S4. Quantitative profiling of SEC fractions for sEVs-CD63-tdT purification.** Raw data for particle number, protein concentration, and tdTomato fluorescence intensity across SEC fractions were analyzed and fitted using Origin (v2025sr1). (A) Particle concentration profile fitted to a single-peak model. The major peak (fractions 6–11, 2.4–4.4 mL) contained >90% of total particles. (B) Protein concentration profile fitted to a multi-peak model. Particle-rich fractions (6–11) accounted for 24.36% of total protein. (C) tdTomato fluorescence intensity fitted to a multi-peak model, with the primary peak co-eluting with particles (fractions 6–11) representing 73.80% of total signal. (D) NTA comparison of sEVs-CD63-tdT and sEVs-Nor, showing no significant difference in size distribution. Data are presented as mean ± SD; n = 3 independent experiments. ns, not significant.

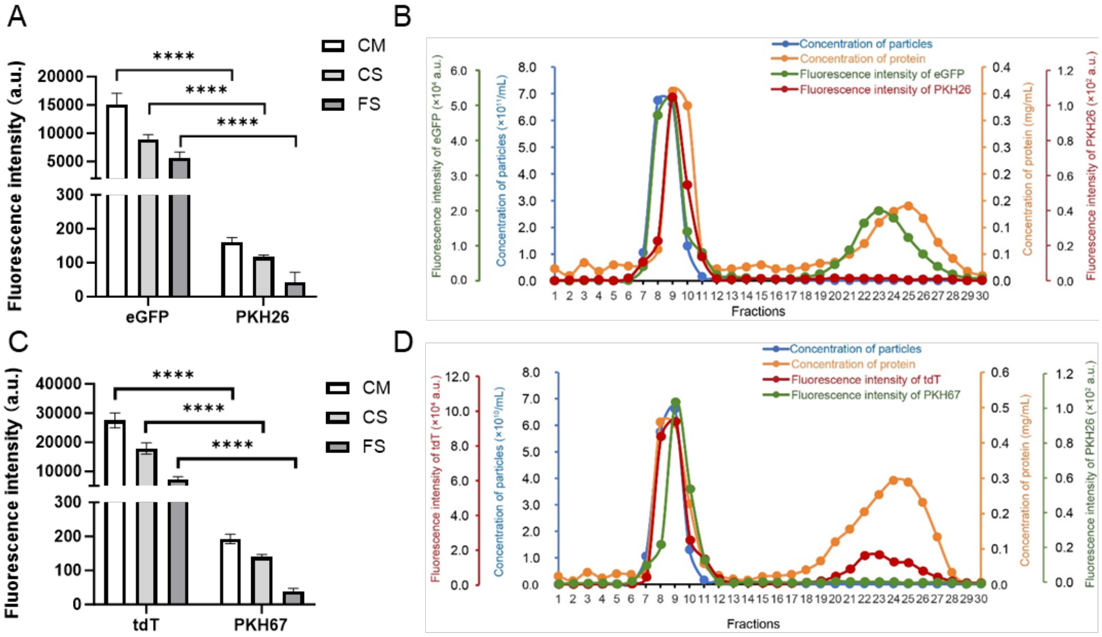


**fig. S5. Purification profiles of PKH26-labeled sEVs-CD63-eGFP and PKH67-labeled sEVs-CD63-tdT.** (A) Volume-normalized fluorescence intensity of PKH26-labeled sEVs-CD63-eGFP in CM, CS, and FS. (B) SEC co-elution profiles of particle concentration, protein concentration, eGFP fluorescence, and PKH26 fluorescence. (C and D) Parallel analysis of PKH67-labeled sEVs-CD63-tdT, showing ultrafiltration efficiency (C) and SEC elution profiles (D). Data are presented as mean ± SD; n = 3 independent experiments. ***$P$ < 0.0001; ns, not significant.

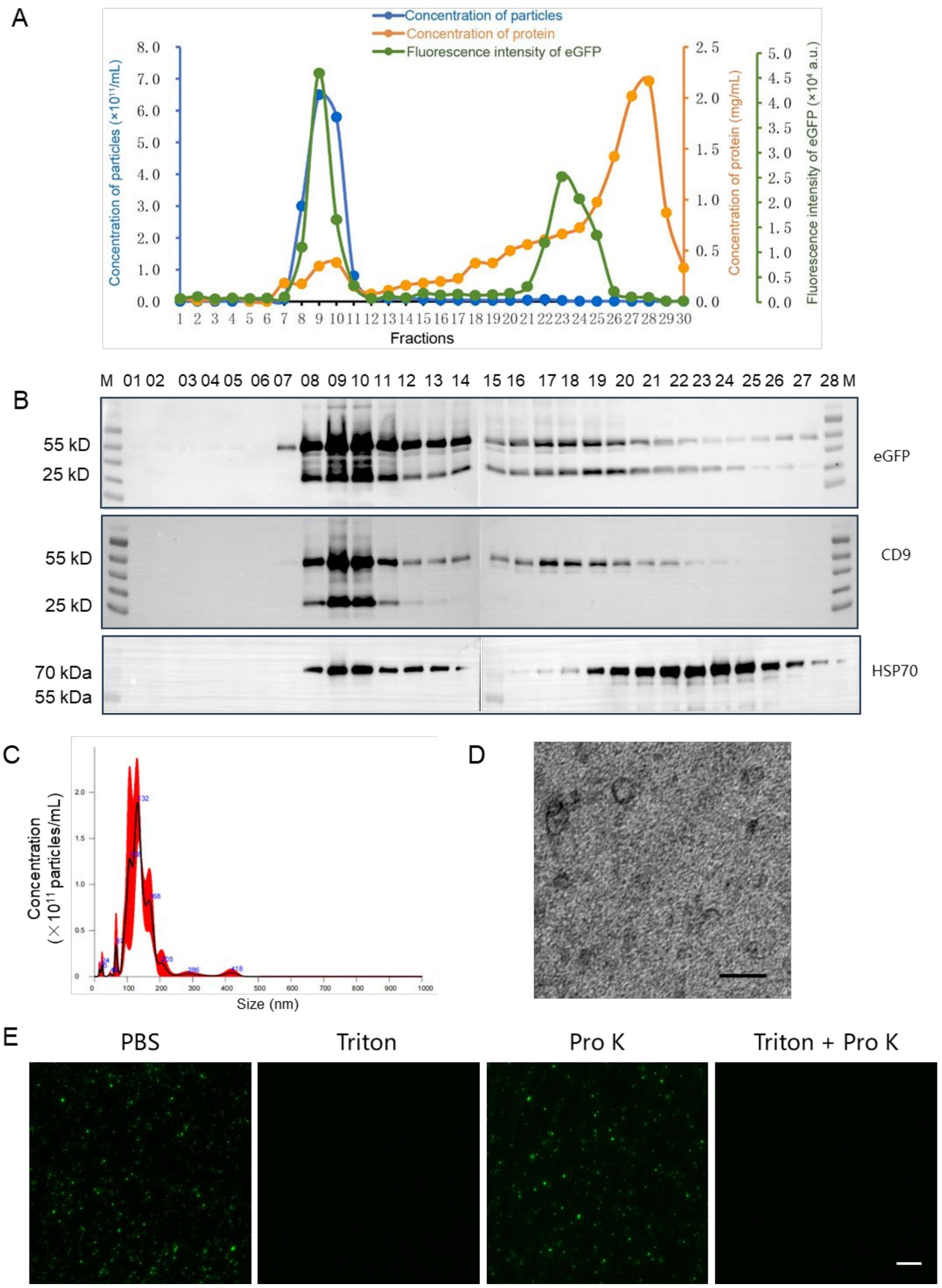


**fig. S6. Generation, isolation, and characterization of sEVs-CD9-eGFP.** sEVs were purified from the conditioned medium of HEK293F cells transiently transfected with pEGFP-C1-CD9 using the REIUS method. (A) SEC co-elution profiles of particle concentration, protein concentration, and eGFP fluorescence. (B) Immunoblot analysis of CD9-eGFP and EV markers (CD9 and HSP70) in SEC fractions. (C) NTA size distribution of pooled fractions 9 and 10. (D) TEM image showing characteristic vesicular morphology. Scale bar, 100 nm. (E) Fluorescence micrographs of pooled fractions 9 and 10 after treatment with Triton X-100, proteinase K, or both, assessing membrane integrity. Scale bar, 10 μm.

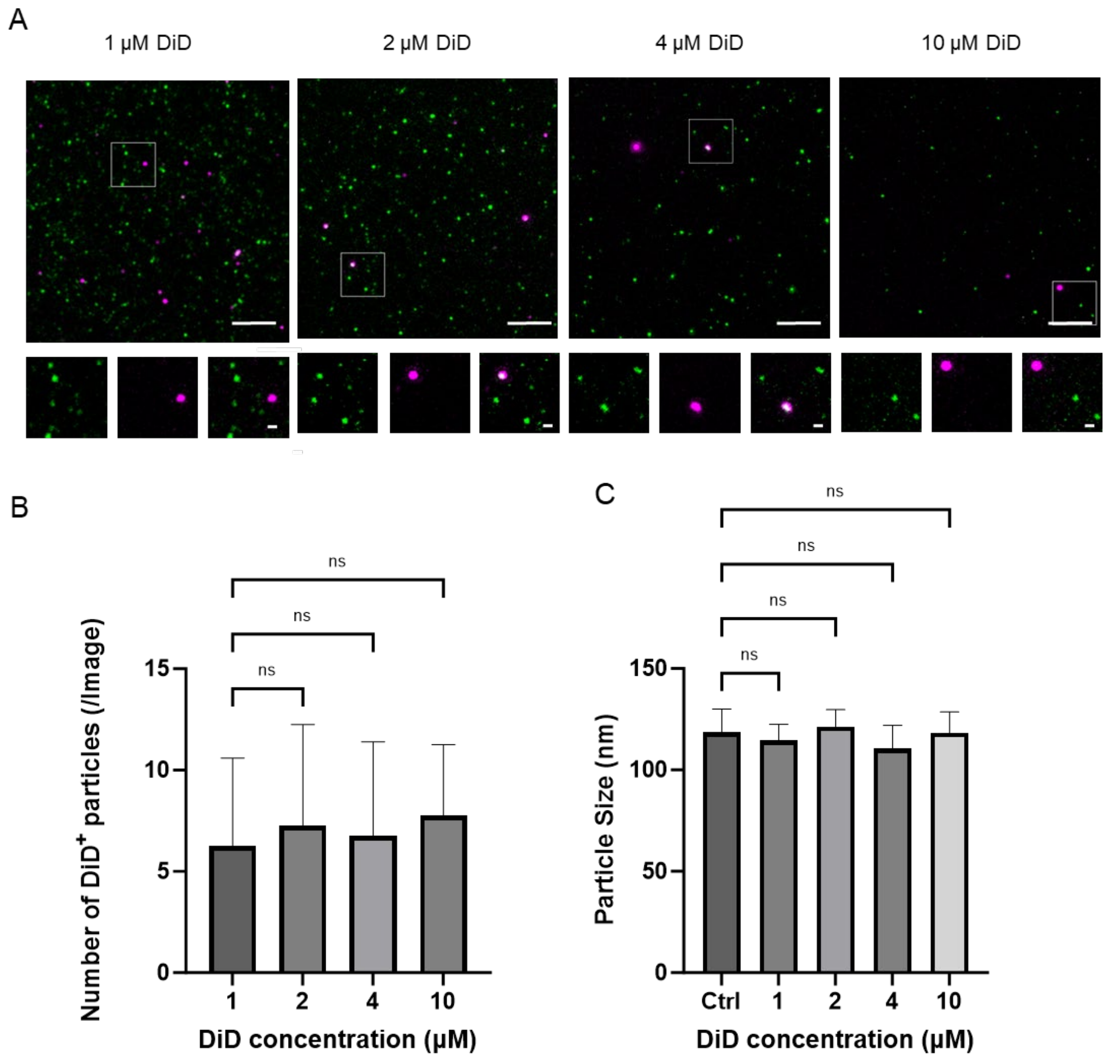


**fig. S7. Concentration-dependent assessment of DiD labeling on sEVs-CD9-eGFP.** (A) Representative fluorescence micrographs of sEVs-CD9-eGFP labeled with increasing concentrations of DiD (1, 2, 4, and 10 μM). (B) Quantitative image analysis of DiD-positive particle counts across the tested concentration range. (C) NTA analysis showing that DiD labeling did not alter the size distribution of sEVs compared with the unlabeled control. Scale bars, 10 μm (overview) and 1 μm (inset). Data are presented as mean ± SD; n = 3 independent experiments. ns, not significant.

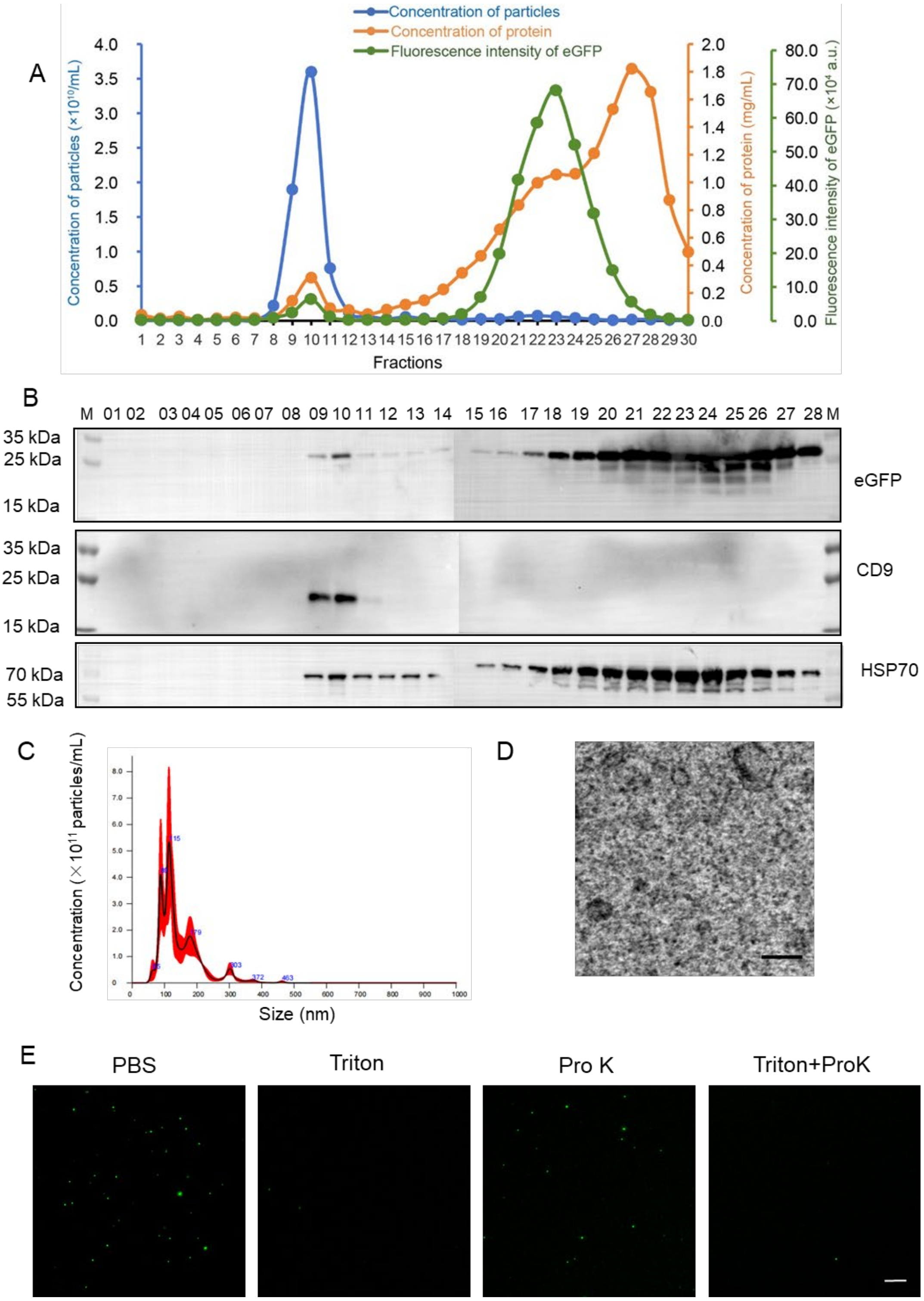


**fig. S8. Generation, isolation, and characterization of sEVs-eGFP.** sEVs were purified from the conditioned medium of HEK293F cells transiently transfected with pEGFP-C1 using the REIUS method. (A) SEC co-elution profiles of particle concentration, protein concentration, and eGFP fluorescence. (B) Immunoblot analysis of EV markers (CD9 and HSP70) in SEC fractions. (C) NTA size distribution of pooled fractions 9 and 10. (D) TEM image showing characteristic vesicular morphology. Scale bar, 100 nm. (E) Fluorescence micrographs of pooled fractions 9 and 10 after treatment with Triton X-100, proteinase K, or both, assessing membrane integrity. Scale bar, 10 μm.

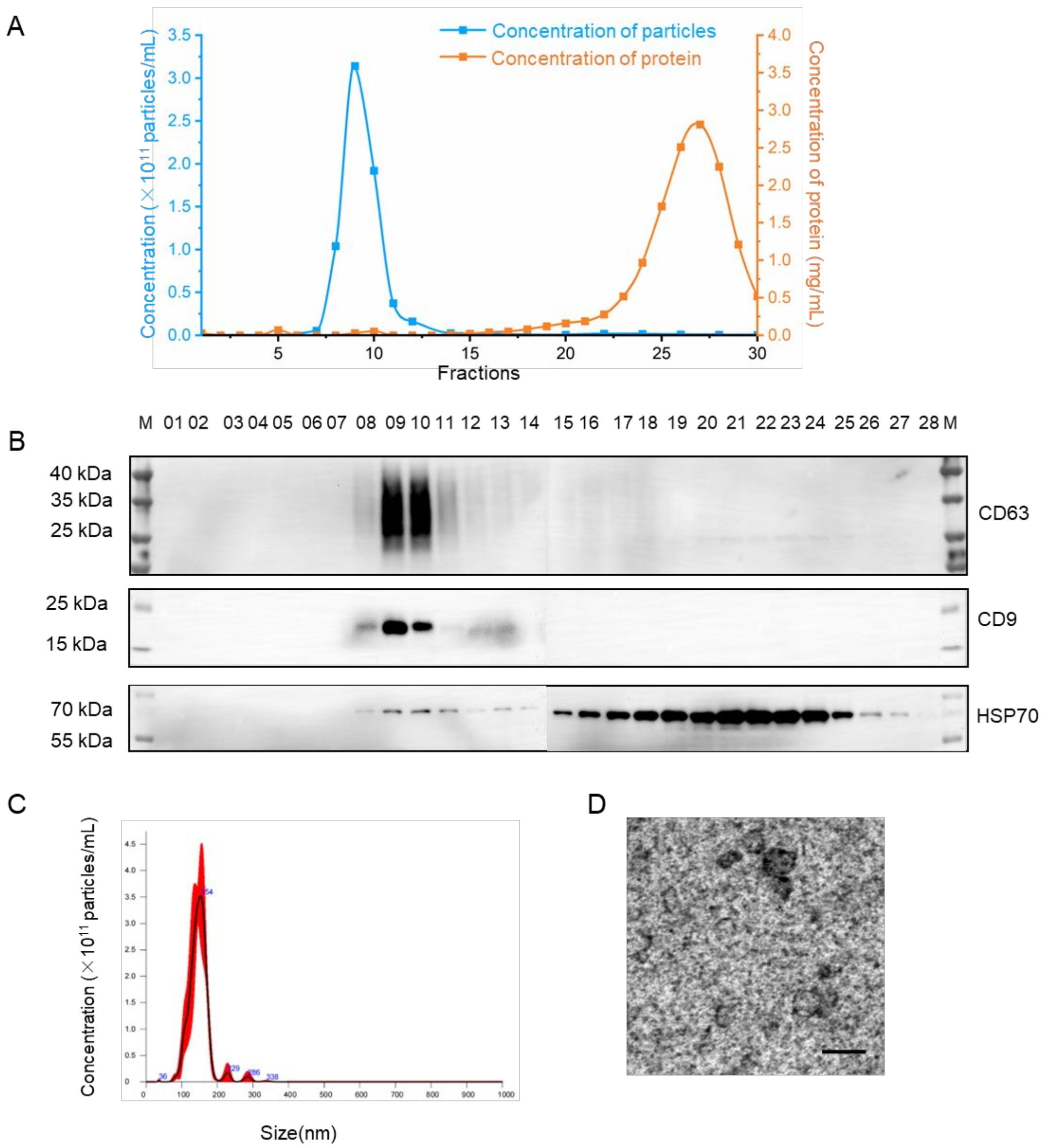


**fig. S9. Isolation and characterization of sEVs-Nor.** sEVs were purified from unmodified HEK293F cells using the REIUS method. (A) SEC co-elution profiles of particle concentration and protein concentration. (B) Immunoblot analysis of EV markers (CD63, CD9, and HSP70) in SEC fractions. (C) NTA size distribution of pooled fractions 9 and 10. (D) TEM image showing characteristic vesicular morphology. Scale bar, 100 nm.

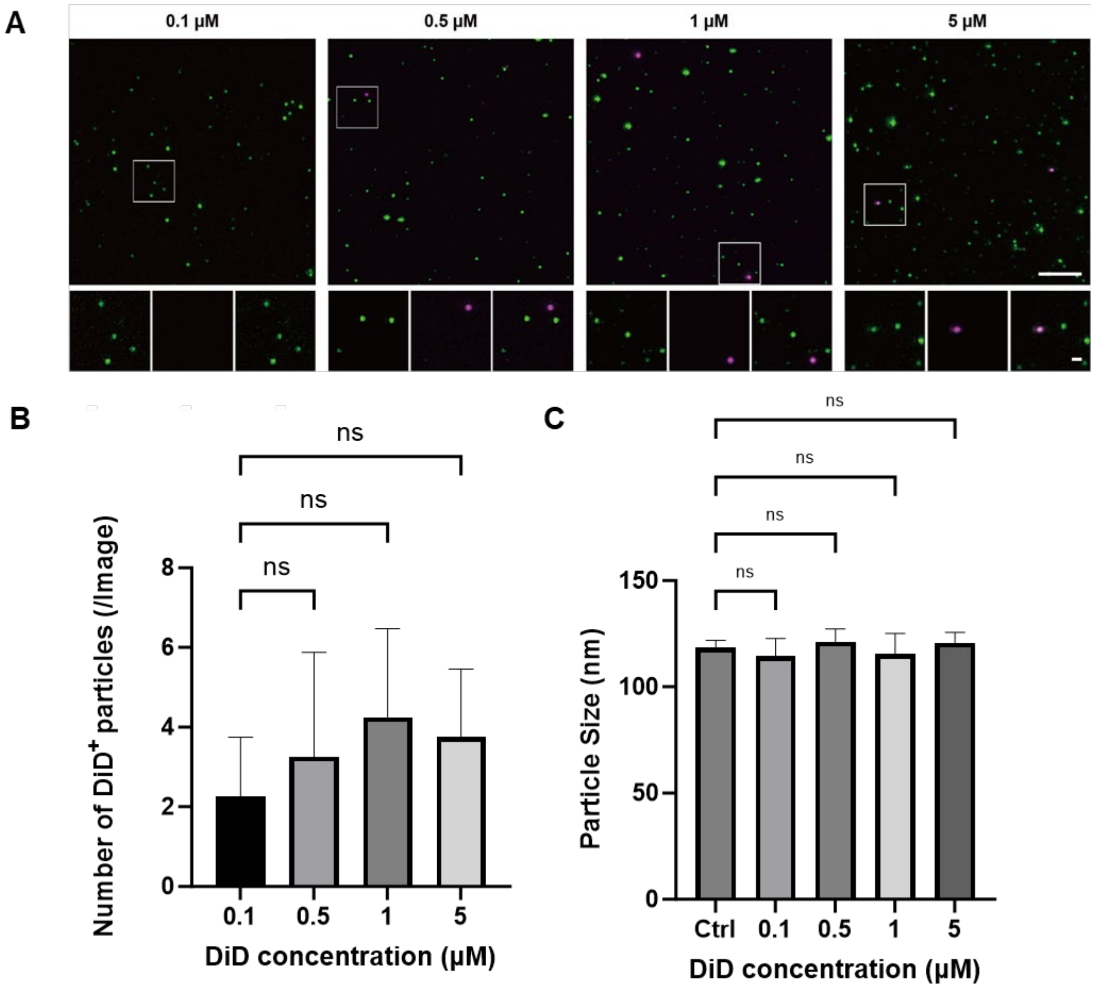


**fig. S10. Concentration-dependent assessment of DiD labeling on sEVs-eGFP.** (A) Representative fluorescence micrographs of sEVs-eGFP labeled with increasing concentrations of DiD (0.1, 0.5, 1, and 5 μM). (B) Quantitative image analysis of DiD-positive particle counts across the tested concentration range. (C) NTA analysis showing that DiD labeling did not alter the size distribution of sEVs compared with the unlabeled control. Scale bars, 10 μm (overview) and 1 μm (inset). Data are presented as mean ± SD; n = 3 independent experiments. ns, not significant.

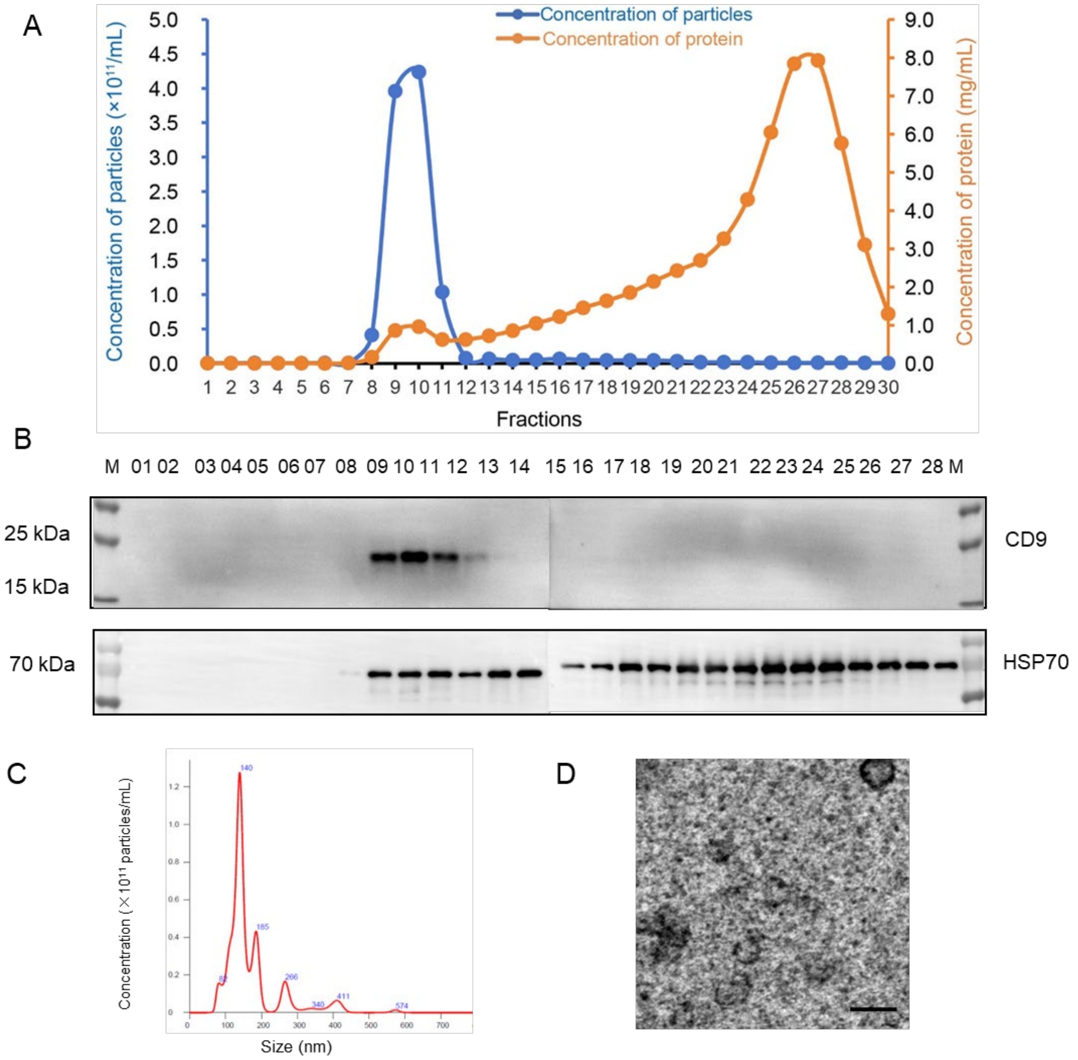


**fig. S11. Isolation and characterization of sEVs derived from CHO cells.** sEVs were purified from CHO cells cultured in chemically defined medium using the REIUS method. (A) SEC co-elution profiles of particle concentration and protein concentration. (B) Immunoblot analysis of EV markers (CD9 and HSP70) in SEC fractions. (C) NTA size distribution of pooled fractions 9 and 10. (D) TEM image showing characteristic vesicular morphology. Scale bar, 100 nm.

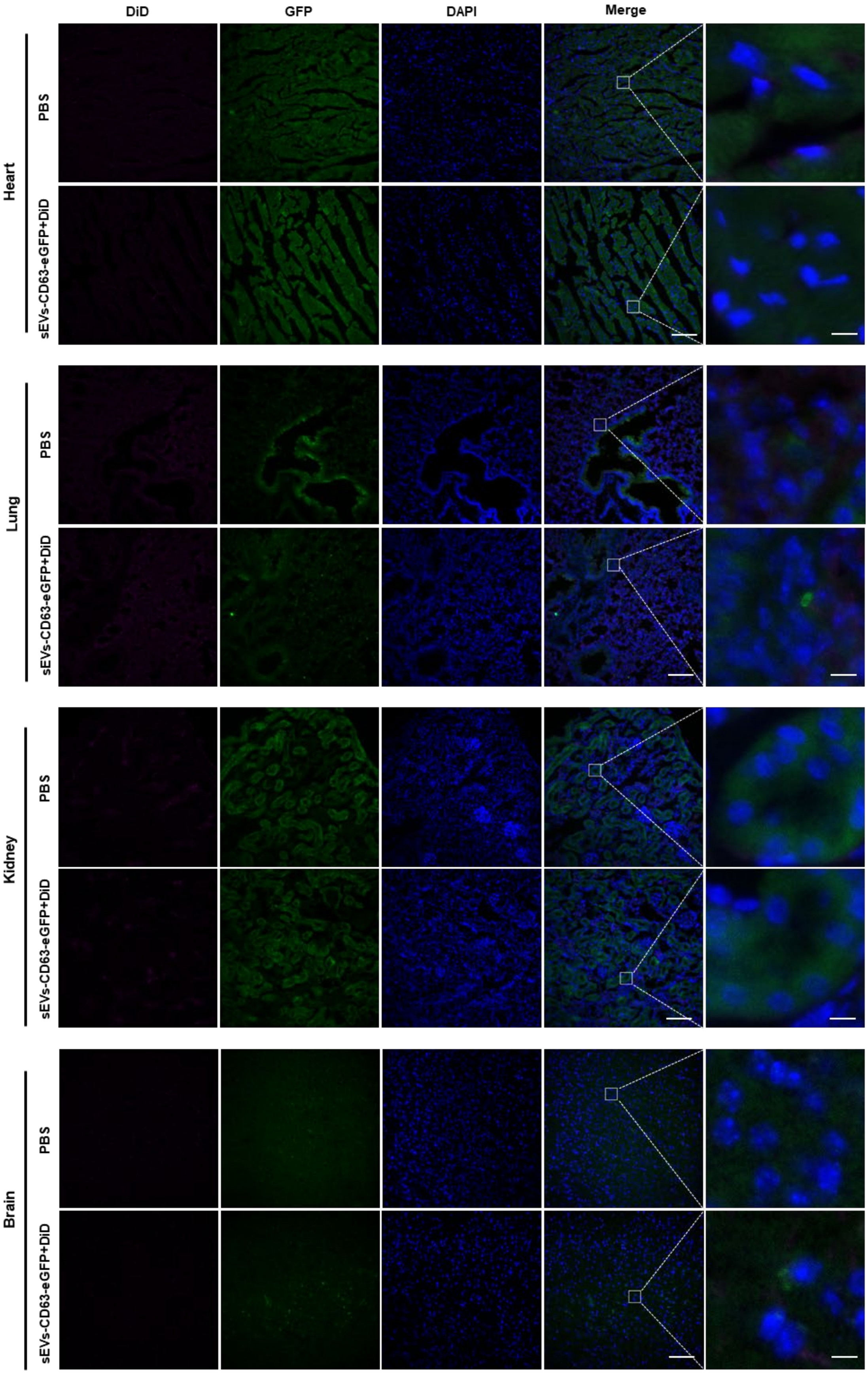
DiD
GFP
DAPI
Merge
Heart
PBS
sEVs-CD63-eGFP+DiD
Lung
PBS
sEVs-CD63-eGFP+DiD
Kidney
PBS
sEVs-CD63-eGFP+DiD
Brain
PBS
sEVs-CD63-eGFP+DiD

**fig. S12. Biodistribution analysis of non-target organs after systemic administration of DiD-labeled sEVs-CD63-eGFP.** Representative immunofluorescence images of heart, lung, kidney, and brain sections from mice 30 min after intravenous injection of REIUS-purified, DiD-labeled sEVs-CD63-eGFP (green, eGFP; magenta, DiD; blue, DAPI), showing no eGFP or DiD signals above background. Control mice received PBS. low-magnification overviews (scale bar, 100 μm); magnified views of the boxed regions (scale bar, 5 μm). n = 3 to 4 mice per group.
